# Real-time intracortical decoding from human motor cortex reveals a task-independent commitment to act that aligns with conscious intent

**DOI:** 10.64898/2026.10.15.759437

**Authors:** Ceci Verbaarschot, Alejandro de Miguel Gomez, Ali Alamri, Jennifer L. Collinger, Charles Greenspon, Aaron Schurger, Robert Gaunt, Uri Maoz

**Affiliations:** Rehab Neural Engineering Labs, University of Pittsburgh, PA, USA; Department of Physical Medicine and Rehabilitation, University of Pittsburgh, PA, USA; Department of Neurological Surgery, University of Texas-Southwestern, TX, USA; Department of Biomedical Engineering, University of Texas-Southwestern, TX, USA; Peter O’Donnell Jr. Brain Institute, TX, USA; Brain Institute, Chapman University, CA, USA; Schmid College of Science and Technology, Chapman University, CA, USA; Department of Neurological Surgery, University of Chicago, IL, USA; Department of Bioengineering, University of Pittsburgh, PA, USA; Crean College of Health and Behavioral Sciences, Chapman University, CA, USA; Anderson School of Management, University of California Los Angeles, CA, USA; Department of Biology and Bioengineering, California Institute of Technology, CA, USA

**Author notes:** Both authors contributed equally to this study (Ceci Verbaarschot, Alejandro de Miguel Gomez).

**Keywords:** Consciousness, intracortical microelectrodes, movement, intention, readiness potential, spinal cord injury, brain-computer interface, volition, agency, brain

## Abstract

Preparatory neural signals that precede movement onset are well established. However, whether they mark a commitment to act, align with conscious intention, or are subject to voluntary control remains under debate. We addressed these questions with a closed-loop experiment: a strategic timing game where a virtual opponent monitored intracortical sensorimotor activity and preempted participants’ actions when their intention to move was detected. In three people with tetraplegia and preserved wrist extension, movement onset was reliably predicted from motor cortex 250-400 ms before action. Although participants attempted to act unpredictably, they were preempted on 94% of trials—and movement output generally followed despite the opponent’s interruption. Participants reported intending to act on 88% of preemptions. The same decoder accurately predicted participants’ actions in a non-competitive task, without retraining. These findings identify a late, task-independent motor-cortical state marking a voluntary commitment to act, which participants report as an intention when interrupted.

## Introduction

Voluntary actions are the result of purposeful decision-making processes that typically culminate in overt movement (Maoz et al., 2019; Mecacci & Haselager, 2015; Verbaarschot, Farquhar, et al., 2019). In contrast to reactive or cued actions, voluntary actions are initiated by a person’s own will to act (Haggard, 2008). As such, voluntary actions are the hallmark of human autonomy and serve as important expressions of accountability and free will (Baumeister, 2008; Hallett, 2007; Mele, 2008; Pockett, 2007; Smith, 2011). The apparent order of events, a thought or feeling of wanting to act followed by the act itself, facilitates the inference of a causal connection between the two (Haggard & Clark, 2003; Wegner, 2017). How the timing and content of these subjective experiences relate to the observed patterns of neural activity that indicate action preparation is not clear.

The preparation for action can be tracked throughout the brain—from the evaluation of sensory input and decision making in prefrontal cortex (Romo & Rossi-Pool, 2020; Rossi-Pool et al., 2020; Soon et al., 2013), to action planning and monitoring in posterior parietal cortex (Aflalo et al., 2022; Klaes et al., 2015; Zhang et al., 2017), and finally preparation of movement execution in motor cortex (Kornhuber & Deecke, 1985; Muller-Putz et al., 2010; Pereira et al., 2018; Shibasaki & Hallett, 2006). On average, these neural preparatory signals for action can already be observed prior to the moment at which a person reports to experience an intention to move. For example, electroencephalographic (EEG) recordings of motor cortex activity show neural activity that is predictive of movement onset up to 500 ms before a person reports an intention to act (Braun et al., 2021; Dominik et al., 2024; Fried et al., 2011; Haggard & Eimer, 1999; Libet et al., 1983; Shibasaki & Hallett, 2006; Trevena & Miller, 2002; Verbaarschot et al., 2015). Similarly, single-neuron activity in posterior parietal cortex has been found to rise significantly above baseline 1000 to 500 ms before intention onset (Aflalo et al., 2022). Functional Magnetic Resonance Imaging (fMRI) recordings show above-chance decoding of decision-specific activity across prefrontal and parietal cortices as early as 10 seconds before that decision entered awareness (Bode et al., 2011; Soon et al., 2008). In contrast to intuitive belief, these studies suggest that the experience of an intention may lag behind the neural preparation for action.

At the single-trial level, the relationship between the onset of early preparatory activity, the experience of intention, and commitment to act remains poorly understood. On the one hand, the observed pre-movement neural activity may mark a genuine commitment to act—a state past which the voluntary action is effectively settled. On the other hand, it may reflect activity leading up to an intention that carries no such commitment, which the actor remains able to override until the movement is underway (Schmidt et al., 2016; Schurger et al., 2012). Determining which interpretation best fits the data has proven difficult because, in traditional experiments, the relative timing of neural preparation for action, intention and movement is reconstructed after the fact. Such experiments can establish that certain patterns in neural activity precede intention reports or action onsets, and can estimate by how much, but they cannot establish whether they signal a commitment to act because the act itself is never placed in jeopardy. For such post hoc analyses, whether the observed preparatory neural activity would still have culminated in movement had something intervened, and whether the actor could have prevented it, never arises as a measurable question.

What is missing is not more precise measurement but intervention: a method that detects the neural antecedent while it is unfolding and acts on it, quickly and accurately enough to preempt the impending movement. Intentions may be decoded during one of three phases: (phase 1) too early when a person is not (yet) aware of their intention to act, (phase 2) on time when a person can confirm their intention to act shortly prior to action, or (phase 3) too late when action execution is already observable (Matsuhashi & Hallett, 2008; Verbaarschot et al., 2016). Deciding whether an early prediction reflects an action commitment that the person cannot yet report, or merely a bias that would have culminated in no action, requires acting on the signal in real-time rather than only recording it for post hoc analysis. Previous attempts to decode motor intentions were either performed offline (Aflalo et al., 2022; Bai et al., 2011; Fried et al., 2011), or lacked sufficient accuracy (true positive prediction rate of ∼60%) before movement onset (Schultze-Kraft et al., 2016; Schultze-Kraft et al., 2021). Although brain-computer interface techniques based on intracortical recordings can achieve fast and accurate decoding of movement intent (Collinger et al., 2013; Dekleva et al., 2021; Struber et al., 2025; Wodlinger et al., 2015), they are conventionally applied to only the final stages of movement preparation and execution (phase 3), where voluntary modulation of motor cortex is well established (Vargas-Irwin et al., 2018). As such, it remains unknown whether the early neural activity (phases 1 and 2) in the region indicative of movement intent is also susceptible to voluntary control. Answering that question requires redirecting the decoded state back onto the actor: BCIs typically use motor-cortical activity to execute an intended movement in the world; here, the neural signal is used to detect an intention while it is still being formed.

In this study, we constructed a real-time, asynchronous classifier (i.e., one that operates continuously rather than only within cue-locked windows) that used multi-unit activity from intracortical microelectrodes in the sensorimotor areas of three people with tetraplegia to decode movement intent. We tested this classifier during a strategic computer game involving an intertemporal-choice task, called “Flip-That-Bucket” (Verbaarschot, Gerrits, et al., 2019). In the game, a bucket was incrementally filled with slime, and participants decided when to flip it over onto the head of a virtual robot opponent via a right-hand wrist extension, contralateral to the implant site. Participants were instructed to try to win the game and were free to use whatever strategy they deemed appropriate. The robot opponent decoded the player’s intention to act from motor cortical activity and tried to flip the bucket before the participant could. This effectively forced participants to play against their own brain. Because the classifier typically fired a few hundred milliseconds before action onset, the game did more than measure the neural antecedent—it intervened on it. On every trial, the participant’s impending action was preempted at precisely the moment their motor-cortical state became decodable. This closed-loop experiment provided the decoded state with behavioral consequences, enabling us to investigate both subjective and neural developments of voluntary movement intent on a single-trial level prior to action onset.

This design lets a single hypothesis—that the detected motor-cortical state marks a general commitment to act—be evaluated through four of its consequences. If the hypothesis holds, the decoded state should: (1) reliably predict imminent action whenever it is left to run its course, which we assessed using silent predictions—logged in the background without intervening in the game; (2) coincide with the participant’s conscious intention to move, which we assessed by asking participants, immediately after each preemption, whether they had wanted to act; (3) resist voluntary concealment and reversal, which we assessed by testing whether motivated participants, who reported trying a range of strategies, could shift their neural activity enough to undermine the classifier; and (4) generalize across behavioral and cognitive contexts, which we assessed by comparing the same movement performed under tightly constrained (a cued reaction-time task) and unconstrained (a self-paced image-scroll task) conditions.

Our results revealed a clear neural signature preceding movement execution, characterized across all three tasks by a rapid increase in neural firing beginning approximately 250 ms before movement onset. Additionally, for voluntary movements only, this shared late increase was preceded by an earlier, more gradual rise in activity that began approximately 500 ms before movement onset. Although participants had intracortical electrode arrays implanted in both motor and somatosensory cortex, movement intent was predominantly visible in motor cortex. Using this signature, a classifier decoded conscious movement intent in real-time with high accuracy, as confirmed by participants’ subjective intention reports. Despite subtle changes in neural activity over time, participants were generally unable to modulate their neural activity sufficiently to effectively conceal their intentions and undermine the real-time classifier. We conclude that, in motor cortex, these neural preparatory signals not only reflect voluntary action commitment but also are accompanied by awareness of it: a state that resists concealment and reliably results in movement.

## Results

Four participants with tetraplegia, who had residual function of wrist extensors, and chronically implanted microelectrode arrays in their motor and somatosensory cortices as part of an ongoing clinical trial (NCT01894802), performed a wrist extension in three tasks with varying levels of volition: (1) a reaction-time task, (2) the Flip-that-Bucket game (hereafter “game task”), and (3) an image-scroll task (Figure 1). During each task, we measured neural activity, muscle activity (electromyography; EMG), and amount of force generated during rest, action preparation, action execution, and post-action. To measure force onset, participants placed their hand in a rectangular measurement device (Figure 2a), with a metal bar right above their hand. Upon wrist extension, a force sensor in the metal bar registered the amount of force exerted on the bar, similar to Dekleva et al. (2024). Force output was monitored in all experimental tasks and was considered an “action” once it exceeded a pre-defined threshold (see STAR Methods).

**Figure 1.**
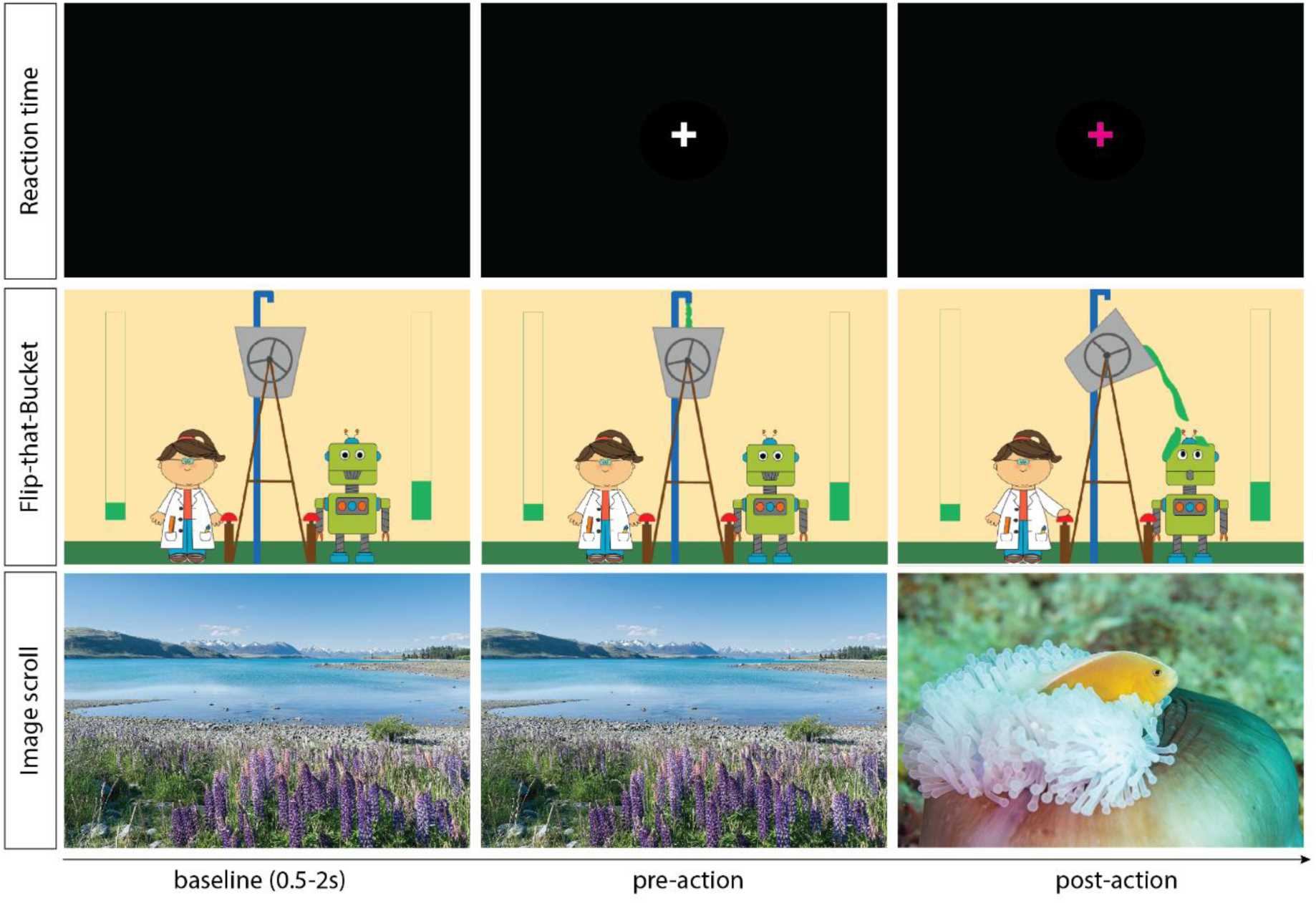
Overview of experimental tasks. (Top) During the reaction-time task, the participant was instructed to perform a wrist extension as soon as a clearly audible tone was presented, after an initial baseline period and a variable wait time (white fixation cross). Once the action was registered by a force sensor, the fixation cross changed color (to purple). (Middle) In the game task, the participant performed the same action in a competitive environment. In each trial, after an initial baseline period, the bucket started to fill with slime. The participant needed to determine the best moment for action to maximize the amount of slime flipped on the opponent, before the opponent flipped the bucket on them. (Bottom) After a short baseline period, the participant was free to scroll to a new image at their own pace. The same action now caused the presentation of a new image.

**Figure 2.**
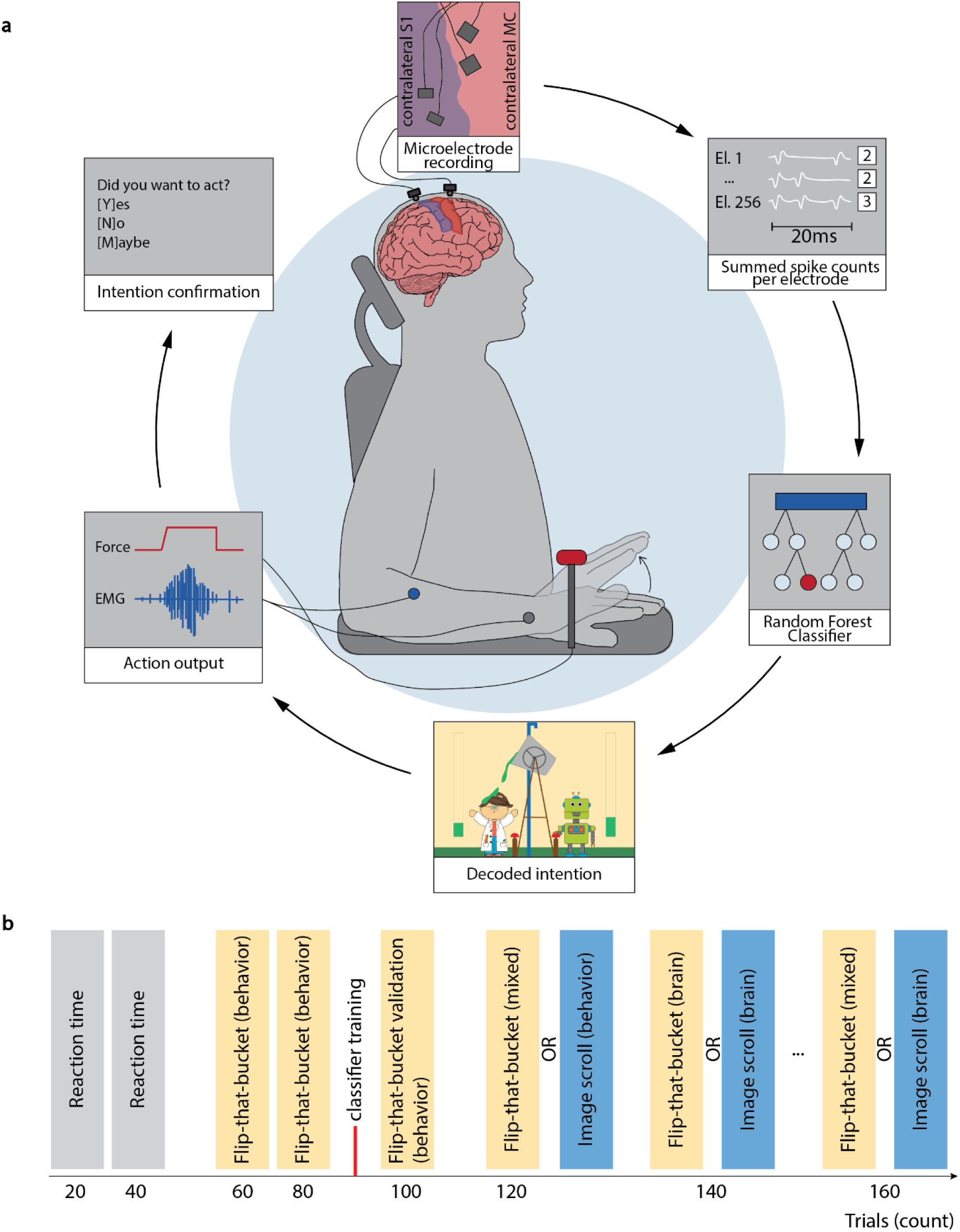
Real-time asynchronous decoding of movement intent and session overview for the game task. (a) Four intracortical microelectrode arrays (Blackrock Neurotech, Inc.) recorded ongoing neural activity in both somatosensory and motor cortex. For each of the 240 (participant P2) or 256 (participants P3, C1) recording microelectrodes, the total number of neural spikes that passed a pre-defined voltage threshold was calculated every 20 ms (see STAR Methods). A random forest classifier analyzed these binned spike counts in a sliding window of 200 ms (10 bins). As soon as the classifier decoded an intention to move, the robot opponent flipped the bucket on the participant’s avatar (i.e., the scientist; Figure 1, middle row), meaning that the participants effectively played against their own brain activity. During the game, both muscle activity and force output corresponding to the participant’s wrist extensions were tracked as ground-truth measures of action onset. Each time the opponent flipped the bucket, the participant was immediately asked to confirm whether they wanted to act at the moment of opponent intervention. (b) Sessions started with two blocks of the reaction-time task (Figure 1, top row), followed by two behavioral blocks of the game task, which were used to train the real-time classifier. The behavioral game blocks were followed by a single validation block of the game. During the validation block, the classifier made silent predictions (recording when it would have flipped the bucket without actually flipping it), providing a chance to measure its performance relative to an uninterrupted stream of participant actions. Additional game or scroll blocks were then tested, depending on participant availability. These additional game blocks were either all controlled by the real-time classifier (brain blocks) or mixed brain and behavior-based prediction (see STAR Methods).

In the reaction-time task, participants were instructed to perform a wrist extension as soon as they heard an auditory cue at an unpredictable moment in time. This task served as a control condition, providing a relative timeline of action-related activity across the three signals we measured (neural preparation, EMG, and force for wrist extension).

In the game task, participants tried to beat a virtual robot opponent in a strategic timing game (Figure 1, middle row). By extending their wrist, they flipped a virtual bucket with slime over their opponent. In “behavioral” blocks of the game, the robot opponent predicted the participants’ force onset by drawing from a Gaussian distribution with a mean and standard deviation calculated from the participants’ movement history (“behavioral predictions”; see STAR Methods). Approximately, 20% of opponent actions would happen 2 s or more before the average participant movement time; 60% would take place 2 to 0 s before the average participant move; and 20% would happen after the participant moved (Verbaarschot, Gerrits, et al., 2019). In “brain” blocks, the opponents’ actions were determined in real-time based on the participants’ ongoing neural activity leading up to the action, meaning that participants effectively played against their own brain (Figure 2a). These “brain-based predictions” were made via a random forest classifier trained to find spiking activity patterns that preceded action onset. Some blocks, termed “mixed” blocks, randomly interleaved behavioral and brain-based predictions. Participants were not explicitly made aware of what triggered their opponent’s actions. Whoever threw the most slime over their opponent in a competitive series of trials won the game.

The image scroll task tracked the same action while imposing no constraints on the action. In this task, the participants scrolled through digital content of their choice (short videos, images, or pages of a virtual comic book) using a self-paced wrist extension. In “behavioral” blocks, the scroll onset was determined based on the participants’ activation of the force sensor. In “brain” blocks, the scroll was performed based on the prediction of movement onset in their ongoing neural activity, using the same classifier from the game task in a zero-shot manner, without additional training. As in the game task, participants were not made aware of what triggered the scroll action.

Experimental sessions comprised blocks of the reaction-time, game, and scroll tasks (Figure 2b). For each participant, we analyzed changes in brain activity and wrist-extension force around action onset across the three tasks. By comparing the force-based registration of action output and the instantaneous increase in EMG activity in response to cue onset in the reaction-time task, we additionally assessed the delay between the actual action output (EMG) and our registration of that output (via force).

We report the experimental results of participants P2, P3 and C1. Due to time constraints, P3 did not perform the scroll task. During multiple sessions, participant C2 was unable to follow experimental instructions, leading to the exclusion of a large part of his data (see STAR Methods). Sessions in which he performed the experimental task as instructed are reported in Figure S1. The results of C2 were similar to those of the other participants in terms of neural activity, force and EMG activity (compare Figures 3 and S1).

**Figure 3.**
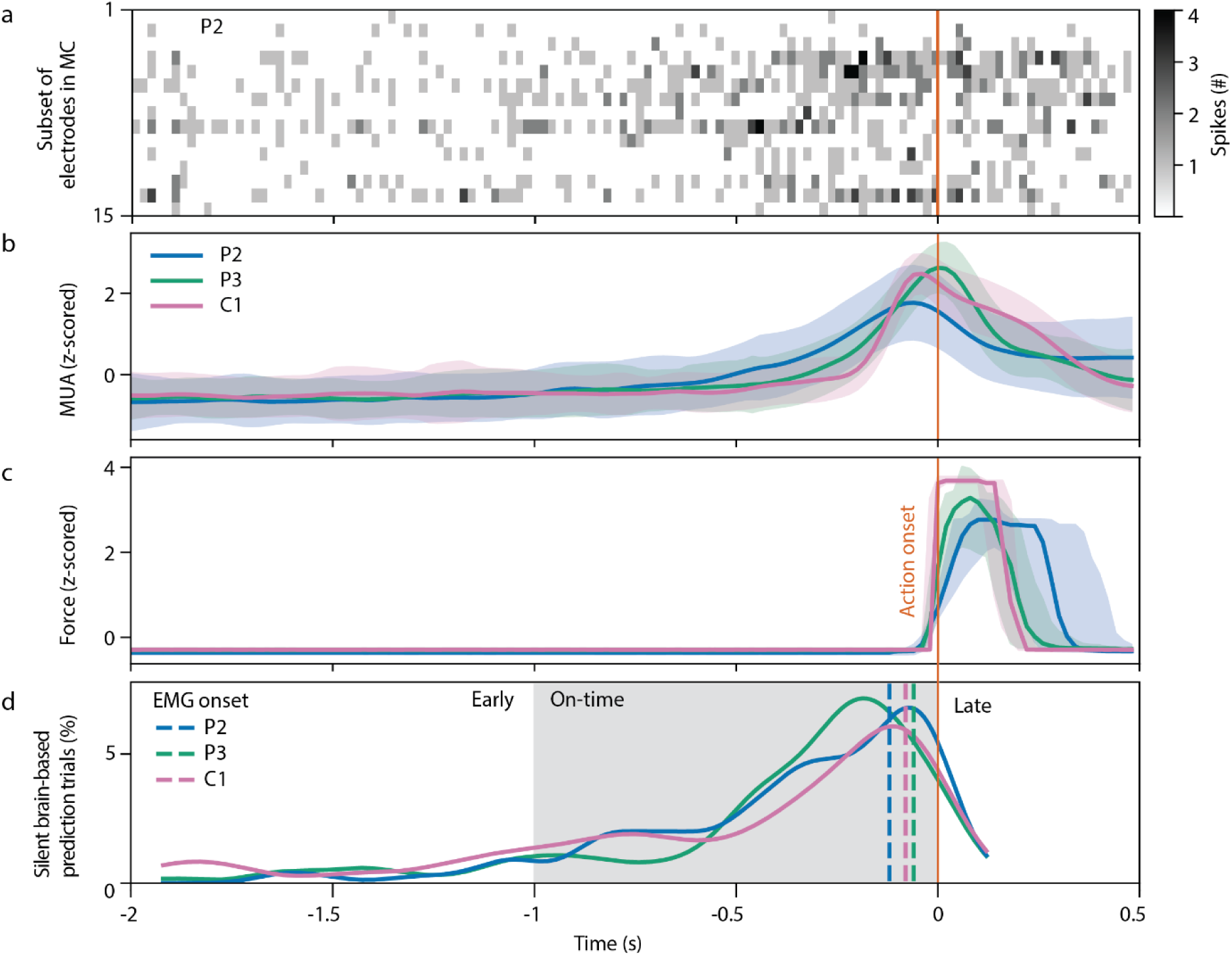
Neural preparation for action and classifier performance during the game task. (a) A typical heatmap (from participant P2), showing the number of recorded neuronal spikes during every 20 ms time bin, from 2000 ms before to 500 ms after action onset (designated time 0) across an example subset of 15 recorded microelectrodes. (b) Mean and standard deviation of the smoothed firing rates across all 240-256 electrodes for the three participants. Firing rates were z-scored per participant and per session. Individual trial results per participant, time-locked to action onset, are provided in Figure S3. (c) Median and 25-75 percentile ranges of the force output recorded for each participant aligned to action onset (time 0). Force trajectories were z-scored per participant and per session. (d) Percentage of silent brain-based classifier predictions aligned to action onset. The time range of on-time predictions is marked in grey. The median onset of muscle activity (EMG onset) is depicted for each participant with a vertical dashed line.

### Decoded neural activity reliably predicts imminent voluntary action

During both the game and scroll tasks, neural decoders generated continuous predictions of the participants’ actions in the background. These “silent predictions” were recorded without triggering the opponent to flip the bucket (game task) or presenting a new image on screen (scroll task). These silent predictions enabled us to assess the timing and accuracy of decoder predictions relative to movement onset while leaving participant behavior uninterrupted. In post hoc analysis, we assessed the timepoint at which the first positive silent prediction occurred per trial. The timing of these silent predictions was then assigned to one of three categories: “early predictions,” happening more than 1 s prior to participant action, as measured through their force output; “on-time predictions,” happening within 1 s prior to participant action; and “late predictions” happening after the participants’ action (Verbaarschot, Gerrits, et al., 2019). Whether these predictions reflect true positives (i.e., they happen when a person experiences an intention to act) cannot be directly determined, as we do not collect intention reports after silent predictions.

Active predictions, which triggered an opponent action (game task), or the presentation of new information (scroll task), were measured in terms of their accuracy based on intention reports. An active prediction was deemed correct if the participant reported an intention to move when asked immediately after the trial ended (Figure 2a).

Our results showed a clear average increase in neural firing starting more than 500 ms prior to action onset (Figure 3a,b). The great majority (84.0%) of silent brain-based predictions happened on-time at 407 ms prior to participant action onset on average (measured relative to force onset) in the validation block of the game task (Table 1, Figure 3c). Throughout the rest of a session (measured in mixed blocks), their occurrence in the game task stayed roughly the same (83.3%) whereas their timing relative to participant action decreased to an average of -244 ms (Table 1). In addition, 14.3% of the silent brain-based predictions happened early, on average 1462 ms prior to action onset in the validation block of the game task, and 1.7% occurred late, around 80 ms after action onset, on average (Table 1). These relative distributions changed in the remainder of the game task (measured in mixed blocks), with 10.7% of early predictions happening around 1596 ms prior to participant action and 6.0% happening late at 154 ms after action onset, on average (Table 1). This decrease in early predictions coincided with a 23.3% decrease in the average wait-time (trial onset to action onset) of participants—from 4.3 s in the validation blocks to 3.3 s in the mixed blocks (Table 1, Figure S2).

**Table 1.**
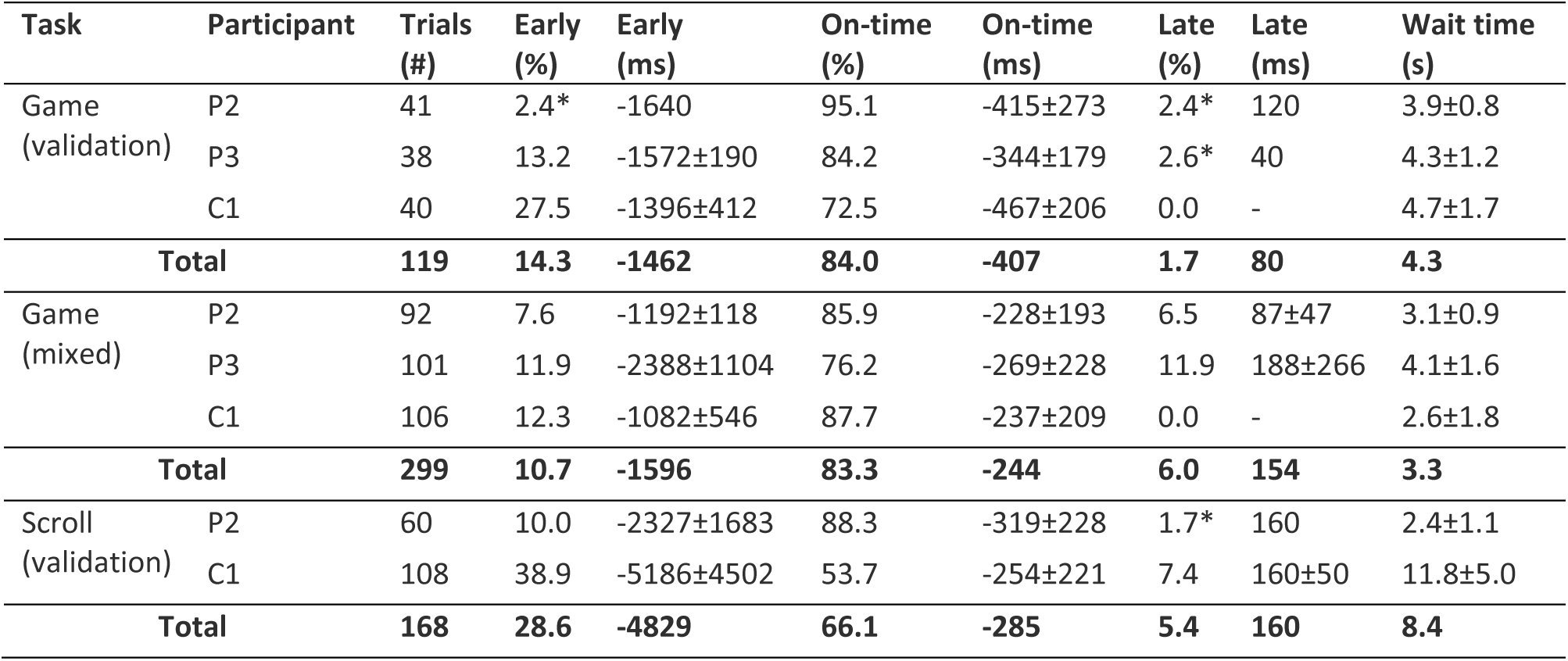
Classifier performance during silent brain-based classifier predictions for the validation and mixed blocks of the game task, as well as the scroll task. For each participant, the table contains the total number of trials, the percentage of trials in each condition, and the mean (± standard deviation) timing of predicted action relative to movement onset. Wait time notes the time between trial onset (or the end of the baseline period; Figure 1) and action onset (in seconds). For all measures, the table also provides the mean across participants (weighted by the number of trials per participant) for each experimental block. *These results comprised a single trial.

| Task | Participant | Trials (#) | Early (%) | Early (ms) | On-time (%) | On-time (ms) | Late (%) | Late (ms) | Wait time (s) |
| --- | --- | --- | --- | --- | --- | --- | --- | --- | --- |
| Game (validation) | P2 | 41 | 2.4* | -1640 | 95.1 | -415±273 | 2.4* | 120 | 3.9±0.8 |
| | P3 | 38 | 13.2 | -1572±190 | 84.2 | -344±179 | 2.6* | 40 | 4.3±1.2 |
| | C1 | 40 | 27.5 | -1396±412 | 72.5 | -467±206 | 0.0 | - | 4.7±1.7 |
|  | **Total** | **119** | **14.3** | **-1462** | **84.0** | **-407** | **1.7** | **80** | **4.3** |
| Game (mixed) | P2 | 92 | 7.6 | -1192±118 | 85.9 | -228±193 | 6.5 | 87±47 | 3.1±0.9 |
| | P3 | 101 | 11.9 | -2388±1104 | 76.2 | -269±228 | 11.9 | 188±266 | 4.1±1.6 |
| | C1 | 106 | 12.3 | -1082±546 | 87.7 | -237±209 | 0.0 | - | 2.6±1.8 |
|  | **Total** | **299** | **10.7** | **-1596** | **83.3** | **-244** | **6.0** | **154** | **3.3** |
| Scroll (validation) | P2 | 60 | 10.0 | -2327±1683 | 88.3 | -319±228 | 1.7* | 160 | 2.4±1.1 |
| | C1 | 108 | 38.9 | -5186±4502 | 53.7 | -254±221 | 7.4 | 160±50 | 11.8±5.0 |
|  | **Total** | **168** | **28.6** | **-4829** | **66.1** | **-285** | **5.4** | **160** | **8.4** |

Most of the silent brain-based predictions, 77.4% across participants, preceded the onset of muscle activation, measured using EMG (Figure S2). On average, muscle activity noticeably increased 87ms prior to the observed increase in force that was used to mark action onset. Specifically, EMG activity increased at −120 ms (interquartile range or IQR = 60 ms) for P2, −60 ms (IQR = 20 ms) for P3, and –80 ms (IQR = 20ms) for C1 (Figure 3d, Figure S2).

Overall, the silent brain-based predictions preceded action onset in 98.3% of trials during the validation blocks of the game task (Table 1, early and on-time combined). Later, during the experimental session, this decreased to 94.0% during the mixed blocks of the game task and to 94.7% during the scroll task (Table 1, early and on-time combined). The classifier accuracy for silent brain-based predictions matched the proportion of trials that the opponent won in active brain-based trials. In blocks where participants played against the brain-based classifier, the opponent won 94.3% of all trials (P2: 93.0% of 100 trials, P3: 86.7% of 60 trials, C1: 99.2% of 120 trials).

Although neural activity was decoded from both motor and somatosensory cortices, offline analyses indicated that the brain-based predictions relied almost exclusively on motor cortex activity (Figures S6, S7). Including somatosensory recordings nevertheless increased overall classifier accuracy for on-time predictions by 2.0 percentage points relative to using motor cortex alone. However, classification based on somatosensory activity alone was far less accurate than classification based on motor cortex activity, with estimated accuracies of 12.3 ± 1.7% and 82.4 ± 0.8%, respectively (Figure S7). The large discrepancy between somatosensory and motor cortex-based decoding accuracies suggests that predictions based on somatosensory cortex alone likely reflect inaccurate and temporary estimates of movement intent rather than reliable and persistent predictions.

Together, these results establish the premise for the rest of the study: once the decoded neural state appears and is left to run its course, movement follows. This state precedes not only the measured force but, on over 75% of trials, the first detectable muscle activity. The neural features supporting its detection arise predominantly from motor cortex.

### Decoder predictions align with conscious intention

In response to active brain-based predictions that triggered opponent bucket flips preempting the participants’ actions, participants reported intending to move in 87.5% of trials (Table 2, brain). In contrast, for behavior-based predictions, which are derived from each participant’s previous wait-time distribution, participants reported an intention to move in only 37.1% of the trials (Table 2, behavior). The wide gap between behavior and brain-based predictions demonstrates the superior accuracy of the latter. This suggests that by the time neural information became decodable in motor cortex (and triggered the bucket flip), participants were aware of their decision to move in the great majority of trials.

**Table 2.** Intention reports after an active behavior-based (top) or brain-based (bottom) prediction in the game task. The behavior-based prediction comprises behavior-only blocks (validation) and behavioral trials during mixed blocks. Similarly, the brain-based predictions comprise brain-only blocks (brain) and brain-based trials during mixed blocks.

| Prediction source | Participant | Trials (#) | Positive intent (%) |
| --- | --- | --- | --- |
| Validation (behavior) | P2 | 32 | 34.4 |
|  | P3 | 22 | 27.3 |
|  | C1 | 19 | 57.9 |
| Mixed (behavior) | P2 | 16 | 31.3 |
|  | P3 | 50 | 40.0 |
|  | C1 | 20 | 30.0 |
| **Total (Behavior)** |  | **159** | **37.1** |
| Mixed (brain) | P2 | 91 | 89.0 |
|  | P3 | 89 | 88.8 |
|  | C1 | 114 | 87.7 |
| Brain | P2 | 93 | 94.0 |
|  | P3 | 52 | 78.5 |
|  | C1 | 119 | 84.0 |
| **Total (Brain)** |  | **558** | **87.5** |

Comparing neural activity preceding behavior- and brain-based active predictions, trials where the participants reported an intention to move revealed a steeper and more prolonged increase in firing for brain-based predictions, exceeding each participant’s baseline firing rate by one z-score unit about 113 ms prior to the opponent’s action, on average (Figure 4a,b top; note that time = 0 is prediction onset). Mean neural activity during the 200 ms preceding prediction onset, corresponding to the neural activity window used by the decoder for classification, was significantly higher for brain-based than behavior-based predictions in all three participants (Welch’s t-tests, Bonferroni-corrected, all p ≤ .018; Table S8). This activity had a long post-prediction tail, reflecting variability in action onset in relation to prediction onset (see Figure 3d). In contrast, for behavior-based predictions, prediction onset occurred in the beginning of a more gradual and generally flatter neural activity increase; however, this profile continued to build and extend through the post-prediction period and into movement execution. As opposed to brain-based predictions, the mean firing rate for behavior-based predictions did not exceed the baseline by one z-score unit until 53 ms after the opponent’s action, on average. Such profile was expected because the behavior-based model was designed to interrupt participants’ actions mostly between 0 and 2 s before they occurred (see STAR Methods). This allowed for much greater variability between predicted and actual intention onset of behavior-based predictions compared to the more accurate brain-based classifier.

**Figure 4.**
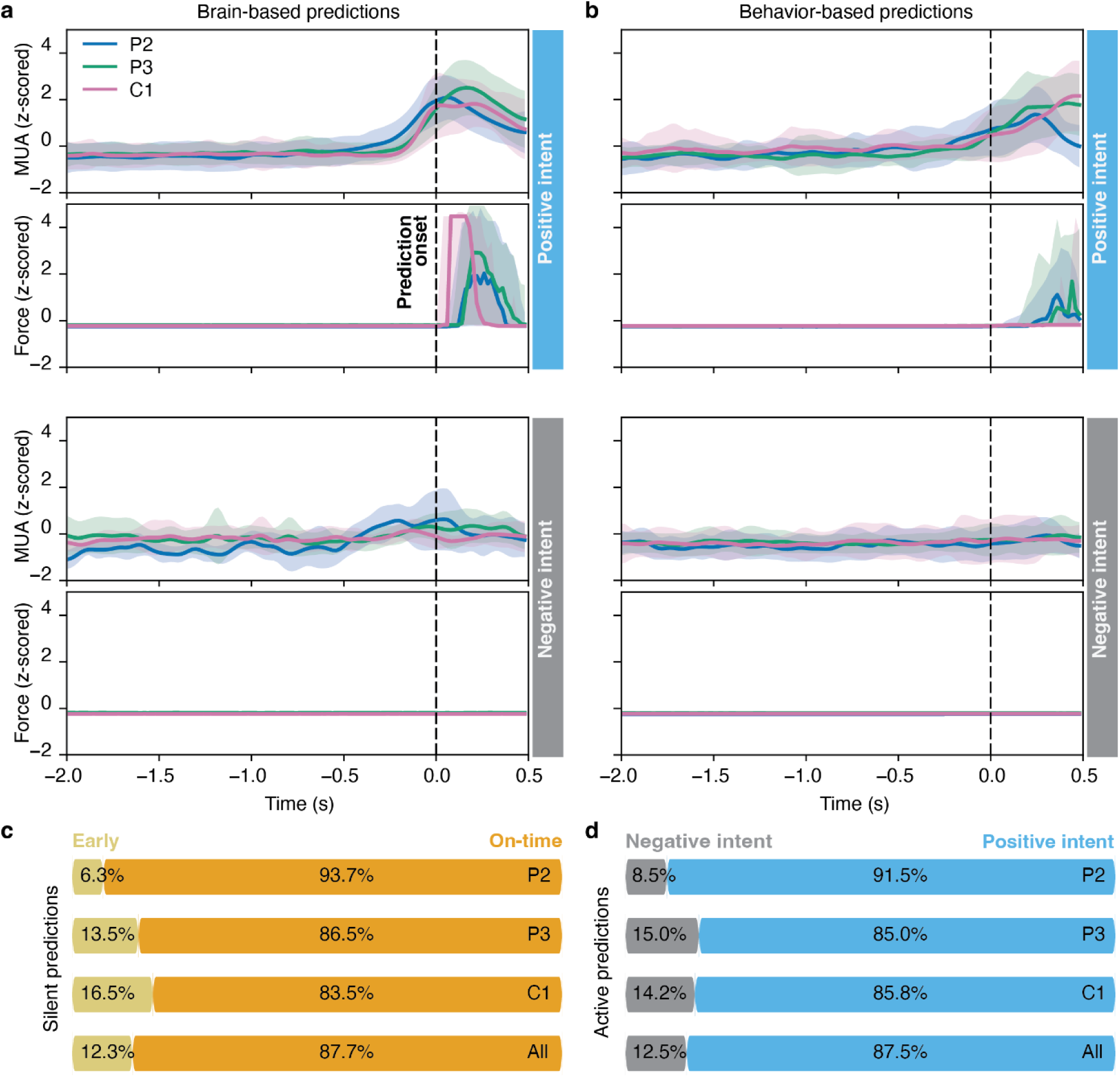
Intention reports vs. classifier predictions during the game task. (a) (Positive intent, top): Z-scored mean and standard deviation of the firing rate prior to active brain-based predictions (time 0) in trials where the participants then reported an intention to move. Z-scoring was done per session and per individual, including both positive and negative intention report trials (see Methods). (Positive intent, bottom): Z-scored median and 25-75 percentile ranges of the force output over the same time range for the same trials. (Negative intent): same as positive intent, but for trials where the participants reported no intention to move after the bucket was flipped on them. Note: In trials with positive intention reports, participant-produced force exceeded the action-triggering threshold after the classifier prediction (time 0) in 92.6% and 67.0% of brain-based and behavior-based predictions, respectively. This indicates that, in most of these trials, the participant acted shortly after the opponent flipped the bucket. (b) Similar to (a), but for behavior-based predictions. (c) Percentage of silent early (<-1 s prior to action onset) vs. on-time (>=-1 s and <0 s prior to action onset) brain-based predictions for each participant and “All” pooled trials. (d): Similar to (c), but for active brain-based predictions—proportion of negative- vs. positive-intent trials.

In both behavior- and brain-based trials, positive-intent predictions were followed by force output exceeding the threshold required to trigger a participant action in 92.6% of brain-based trials and 67.0% of behavior-based trials (Figure 4a,b top). In other words: participants carried out the wrist extension, causing an increase in measured force, even though they already lost that trial (the opponent flipped the bucket first). As a result, participants frequently accused the opponent of “cheating,” or stated that it felt like they were dumping the slime on top of themselves instead of the opponent—e.g., [C1] “I am the robot, not the scientist” (Video S4, S5).

We repeated the same analysis for the trials where the participants reported no intention to act after the opponent flipped the bucket on them (negative-intent trials; Figure 4a,b bottom). Pre-onset neural activity was also significantly higher for brain-based than behavior-based predictions in all three participants (Welch’s t-tests, Bonferroni-corrected, all p ≤ .025; Table S8). This difference is not unexpected, since classifier predictions were necessarily triggered by detecting spiking activity patterns different than baseline, even if participants subsequently reported no intention to act. In either case, the absence of an intention to act was coupled with an absence of increased force output, with force exceeding the threshold required to trigger a participant action in only 3.6% of brain-based trials and 1.6% of behavior-based trials, meaning that—unlike the positive-intent trials—participants did not generally move after the onset of the prediction.

The proportion of negative vs. positive intention reports associated with active classifier predictions closely matched the distribution of early vs. on-time silent predictions (Figure 4c, d). Considering only trials in which silent brain-based predictions preceded participant action (excluding the relatively uncommon late predictions, which would have never interrupted the participant had these been active; see Table 1), predictions happened early in 12.3% of cases, and on-time in 87.7% of cases, on average. This closely mirrored the mean proportion of active brain-based predictions followed by positive (87.5%) or negative (12.5%) intention reports (Figure 4c,d). This correspondence persisted for individual participants as well as on average. These results suggest a time-dependent relationship between brain-based predictions and intention reports, with early and on-time predictions corresponding to negative and positive intention reports, respectively.

Importantly, this correspondence was also evident for behavior-based interventions. By design, the behavior-based model did not generate or log continuous outputs; instead, it determined when to intervene using a Gaussian distribution derived from participants’ wait times. The theoretical distribution predicts that 50%, 30%, and 20% of interventions should occur early, on-time, and too late, respectively (see STAR Methods). After excluding the too late proportion and renormalizing the remaining percentages, as was done for brain-based predictions, the expected distribution is 62.5% early and 37.5% on-time. This closely matched the proportion of negative and positive intention reports elicited by behavior-based interventions, which were 62.9% and 37.1%, respectively.

Overall, the task-related activity we decoded in motor cortex was closely aligned in time with participants’ reported intention to move. The decoded state was therefore not simply a covert precursor to which the actor had no access: on the great majority of trials, it emerged together with, or just before, the reported intention, while the minority of predictions that occurred too early was matched —in both proportion and timing—by the minority of trials on which participants denied having wanted to act.

### Changes in neural activity during gameplay reflect signal drift rather than strategic modulation

Despite the participants’ efforts to act unpredictably and win the game, they were only rarely able to beat the brain-based opponent during the game task (Video S4, S5). Participants were visibly engaged: they argued with the opponent, accused it of cheating, and returned game after game with new tactics. Participant C1, for example, alternated between moving as quickly as he could and waiting far longer than usual. Participant P3 reported strategies like: “I try to beat the robot by thinking about other stuff, like singing a song. I try to make my mind think of something else. Or like, I don’t want to act, and then super spontaneously flip [the bucket].” This suggests that they were striving to find strategies to win against the real-time classifier.

Although it remains unclear whether such strategies could actively reshape the neural signature of intent, we did observe a shift in the firing rates leading up to movement onset from the first to the last trial in each experimental session. Across all three participants, subsets of electrodes exhibited either a gradual linear increase or decrease in firing rate over the course of the session, effectively altering their baseline activity (Figure 5a,b,c). To quantify this effect, we fit linear models to each electrode’s firing rate across all trials within a single session, using the fitted slope as an estimate of the rate of change in activity over the session, to later cluster them into groups of electrodes that provided evidence of consistent increases (“positive” cluster) or decreases (“negative” cluster) in firing rate (see STAR Methods). Notably, these electrode groups—identified during the pre-movement period only—generalized to a baseline period 1 s before trial onset (Figure S9), revealing the same offset in activity during a period in which concealing one’s intention would presumably be least necessary. This generalization to baseline suggests that the activity offset reflects a task-independent neural drift, rather than a process specifically tied to intention or its strategic concealment.

**Figure 5.**
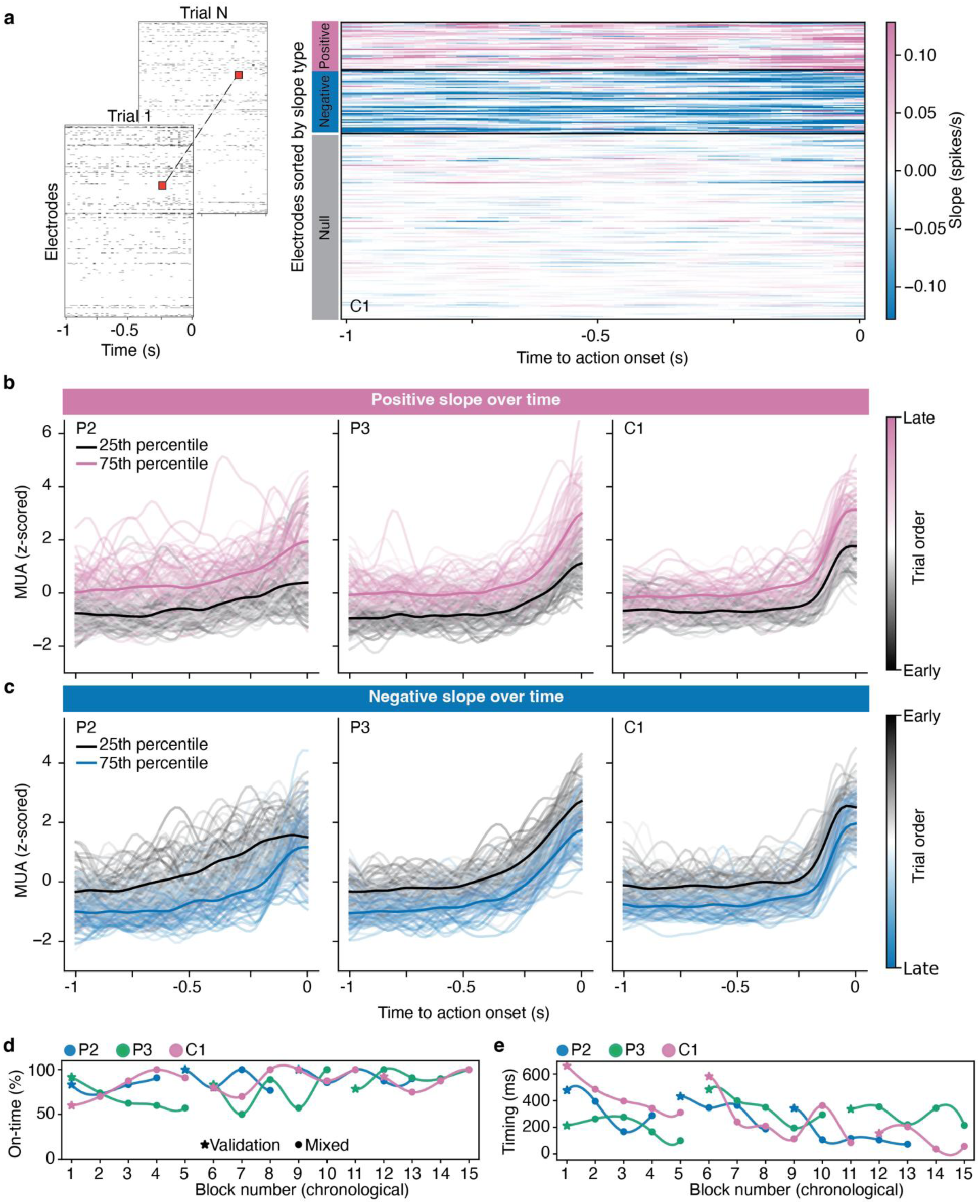
Neural modulation over time. (a) Example data from a typical session (participant P3). Using a binomial test statistic, the changes in pre-movement firing rates over the time course of each experimental session were calculated per electrode across the last second before action onset (see STAR Methods). Based on the outcome of the test, electrodes were grouped as having a significant increase / decrease in firing rate over time (positive / negative) or neither (null). (b) Progression of firing rates across electrodes that show a significant increase in their firing rates over time. The thick lines show the average activity across all positive electrodes in the 25th and 75th percentiles, when the trials were sorted temporally, from first (earliest) to last (late) in each session (across all the blocks in that session). Each thin line shows the pre-movement firing rate on a single trial, colored by the time of occurrence within a session. (c) Same as (b) but for electrodes with a negative slope in their firing rate. (d) Mean percentage of on-time brain-based classifier predictions across multiple Flip-that-Bucket games and across sessions, ordered chronologically with line breaks between sessions. Each line is color coded for each participant. The overall percentage of on-time predictions increased moderately but significantly over time across all participants (Table S11). (e) Mean timing of the on-time brain-based classifier predictions across sessions, ordered chronologically. Lines are color coded for each participant. As a time progresses, the timing of opponent predictions gets significantly closer to participants action onset (P2 and C1 only, not significant for P3, see Table S11).

The gradual changes in firing rate nevertheless shifted the neural activity distribution away from the original training data, resulting in a growing mismatch between the statistics on which the real-time system was trained and those encountered during decoding as the session unfolded. Importantly, electrodes exhibiting systematic increases or decreases in firing rate accounted for nearly 70% of the model’s total feature importance (with “positive” and “negative” clusters contributing approximately 50 and 20%, respectively), despite comprising only 37% of all recorded channels (Table S10). Moreover, nearly 80% of these drifting channels were recorded from the motor cortex arrays. In other words, because most of the model’s predictive power relied on a small subset of motor channels whose firing statistics drifted over the session, these distributional changes were likely to have decreased real-time decoding performance. This may explain the session-wide changes in silent prediction statistics observed across validation and later mixed blocks (Table 1), even though the drift itself did not appear to be intention-specific.

Despite the measured changes over time, participants were still unable to beat the brain-based opponent. Across participants and recording sessions separated by days to months, the percentage of on-time predictions increased modestly as the game task progressed (OLS Linear regression, β = 1.47 percentage points per block, p = 0.003), with no evidence that this trend differed between participants (block x participant interaction, nested-model F-test: F(2, 37) = 0.46, p = 0.636; Table S11). In contrast, the timing of silent on-time predictions relative to participant action changed in a participant-dependent manner (block x participant interaction, nested-model F-test: F(2, 37) = 10.92, p < 0.001; Table S11), moving closer to action onset over time in P2 and C1 but not P3. Participants’ wait times also decreased over time (Table 1), though this had no measurable effect on their success rate at beating the opponent.

Taken together, these findings indicate that sustained and inventive effort did not enable participants to meaningfully conceal or reshape the motor-cortical activity on which the decoder relied. The changes we observed were gradual, likely task-independent, and present even at rest, making them more consistent with drift in the recorded neural population or with fatigue than with strategic control. For practical purposes, these changes were not sufficient to alter the outcome of the game.

### The decoded neural state generalizes across distinct behavioral and cognitive contexts

Interestingly, although the required action was identical across tasks—an extension of the wrist— the timing and amplitude of the average firing rate prior to and during the action differed across tasks (Kruskal–Wallis, all p < 0.001: Figure 6a). Cue-based (reaction-time) actions showed the steepest and highest increase in neural firing prior to action onset. This was confirmed by significantly higher peak pre-movement MUA compared to all other tasks across participants (all Bonferroni corrected p < 0.05, Mann Whitney U; Figure 6b, see Table S12-13 for full statistics). Furthermore, the normalized neural profiles of these trials exhibited significantly lower area-under-the-curve (AUC; see STAR Methods) values compared to other tasks (reaction-time vs game task, P2 and P3, Bonferroni corrected p < 0.0001; and reaction-time vs image scroll, P2 and C1, Bonferroni corrected p < 0.0001, Figure 6c, see Table S14-15 for full statistics). In this context, AUC refers to the area under each trial’s min-max normalized MUA activity profile during a 1-second pre-movement window, capturing both how early activity rises and how long it remains elevated independent of absolute activity magnitude and instead emphasizing the shape of the temporal profile. Lower AUC values therefore indicate that activity stays low for most of the window and increases only shortly before movement onset, consistent with a more abrupt, “steep” rise.

**Figure 6.**
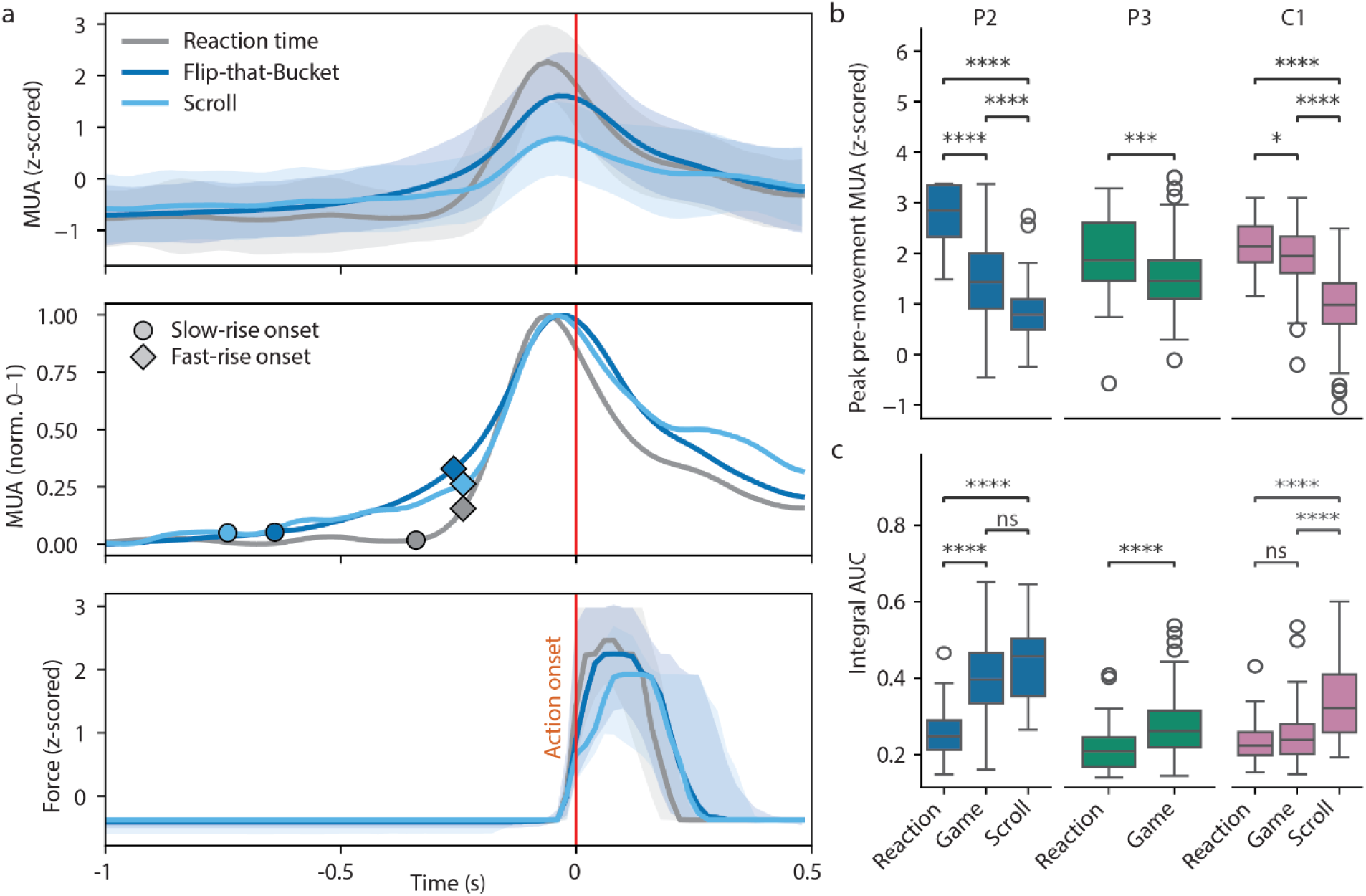
Neural differences between tasks. (a) Top: Mean Gaussian-smoothed firing rate across participants prior to participant action (time 0) for the reaction-time task, game task, and image-scroll task. Firing rates were smoothed using a Gaussian kernel spanning ±160 ms around each time point (σ = 40 ms), with the time axis aligned to the center of the smoothing kernel. A clear difference in neural firing can be observed across tasks, with the reaction-time task having the steepest increase in neural firing prior to action onset and the image scroll the lowest. Firing rates were z-scored per participant and per session, preserving the differences across tasks. Middle: Same as Top panel, but showing the min-max normalized firing rate trace across participants to remove differences in scale and highlight their shape. A clear order of increased firing rates can be detected across tasks, with the game and scroll tasks having the earliest onsets of rising activity, and the reaction-time task the latest. Circles and diamonds mark the slow-rise and fast-rise onsets identified by the segmented-regression breakpoint analysis; the fast-rise onset is closely aligned across all three tasks at approximately 250 ms before action onset (see STAR Methods for details about the implementation). Bottom: Mean force output across participants, for each task. Similar to Top panel; force trajectories were z-scored per participant and per session. (b) Difference in peak pre-movement spiking activity across experimental tasks, separated into participants. Significance codes: * p < 0.05, ** p < 0.01, *** p < 0.001, **** p < 0.0001; otherwise, non-significant (ns). (c) Difference in area under the curve (AUC) of the normalized pre-movement spiking activity per participant across experimental tasks. Significance markers are identical to (b).

Competitive (Flip-that-Bucket) actions showed an earlier and intermediate increase in firing magnitude. Peak MUA for the game task was significantly lower than reaction-time trials (all Bonferroni corrected p < 0.05), but significantly higher than scroll trials (Bonferroni correct p < 0.0001 for C1 and P2). Finally, completely unrestricted (image scroll) actions showed an equally early yet smaller increase in neural firing, characterized by the lowest peak MUA across all conditions (all pairwise comparisons with Bonferroni corrected p < 0.0001) and significantly higher AUC values than reaction-time trials, consistent with an earlier, more gradual ramp-up of activity. These differences in neural preparation were accompanied by differences in mean force output during the first 200 ms after action onset (Welch’s ANOVA, p < 0.001; see Tables S16-17 for full statistics of the force analysis; Figure 6a, bottom row). Mean z-scored force was lower during the unrestricted task (1.39) than during either the cue-based (1.76) or competitive (1.77) task (Bonferroni-corrected p < 0.001 for both comparisons), whereas force did not differ between the cue-based and competitive tasks (Bonferroni-corrected p = 1.00).

To determine whether these differences in neural profile shape reflected distinct phases of pre-movement activity, we fitted a two-breakpoint segmented regression independently to the trial-averaged, min–max-normalized MUA profile for each task (Figure 6a, middle; see also STAR Methods for implementation details). This analysis identified the onset of an initial slow rise and a subsequent fast rise. In both self-paced voluntary tasks, activity departed from baseline early and gradually: the slow rise began 640 ms before movement onset in the game task and 740 ms before movement onset in the image-scroll task. By contrast, the onset of the fast rise was closely aligned across all three tasks, occurring 260 ms before movement in the game task and 240 ms before movement in both the image-scroll and reaction-time tasks. Because each task was fitted independently, this convergence points to a stereotyped final build-up of neural activity that is largely independent of behavioral context and locked to movement onset at approximately 250 ms before action. Although the model also identified an apparent slow-rise onset in the reaction-time task at 320 ms before movement, its proximity to the fast-rise onset at 240 ms leaves little evidence for a genuinely distinct slow phase. This result is consistent with the later and steeper pre-movement increase observed for cue-based actions, which do not require the same neural preparation of an internally generated movement intention.

Despite task-related differences in peak neural activity and cognitive demand, the very same classifier trained on the game task also performed well in the scroll task. During the scroll task, the brain-based classifier accurately predicted 98.3% of P2’s actions and 92.6% of C1’s actions prior to action onset (Table 1, scroll task, early and on-time measurements combined). Although active predictions in this task were not followed by the intention question used in the game to assess prediction accuracy, spontaneous quotes from both participants during the task suggest these predictions were in line with their intention to act (Video S18, S19): [P2] “I could just sit back and not do anything and it will scroll for me”, [P2] “There was a longer [wait time] and it stayed until I actually intended to [act]”, [C1] “[it] proceeded right as I was about to move, like it beat me to the drop”, [C1] “it’s like it is reading my mind”, [C1] “It was definitely doing whatever I wanted.”, [C1] “I can just twitch [my wrist] and it will move. It is like I am barely even flexing [my hand] and it moves. That is cool… It felt like I only had to think about going to the next slide and it was going.”

During the scroll task, the percentage of on-time brain-based classifier predictions was similar to the game task for participant P2 (88.3%), but decreased for participant C1 (53.7%, Table 1). Compared to the game task, C1 clearly increased his wait-time in the scroll task (11.8 s vs. 4.7, on average), which may explain the large increase in early predictions (38.9%, Table 1) happening 5186 ms before action onset on average, at the expense of on-time predictions. These early predictions could reflect false positives, or an urge rather than a commitment to act.

The activity we decoded was therefore not a product of the competitive setting. Its amplitude and time course varied substantially with behavioral and cognitive context—steep and late when the action was cued, early and shallow when it was entirely self-paced. Yet a single decoder, trained on the competitive game task and applied without modification, predicted action equally well in a task the participants performed purely for their own entertainment.

## Discussion

Across three tasks—cued, competitive, and entirely self-paced—participants produced the same voluntary wrist extension. In the latter two tasks only the decision of when to act was left up to the participants (Brass & Haggard, 2008). Given the nature of these two tasks, we expected the interval between the reported onset of the intention to move and action execution to be around 200 ms, as in Libet-type paradigms (Libet et al., 1983; Verbaarschot et al., 2025). Broadly, that was what we found. Although motor-cortical activity began to rise a few hundred milliseconds earlier, movement intent did not become reliably detectable until 400–250 ms before action—and by that point, participants almost always reported having intended to act. Below we take up in turn the four consequences of the hypothesis that this decoded state marks a neural commitment to act: that it predicts imminent action, that it coincides with conscious intention, that it resists concealment and reversal, and that it generalizes across diverse cognitive contexts.

Movement preparation is widely observed before movement execution, whether imagined, attempted or actual (Churchland & Shenoy, 2024; Georgopoulos et al., 1989; Rigotti-Thompson et al., 2026; Vargas-Irwin et al., 2018). In line with the relative timing reported in previous research (Fried et al., 2011; Libet et al., 1983), we found a slow increase in neural firing beginning about 500 ms before action onset, followed by a sharper, faster rise at approximately 250 ms, across microelectrode arrays in the motor and somatosensory cortices of three tetraplegic participants during self-initiated, voluntary actions (Figures 3 and 6a). Based on their motor cortex activity, we could asynchronously detect 94% of the participant’s intentions in the game task (Table 1, mixed). At the same time, in 92.6% of trials when we intervened and flipped the bucket on the participants’ avatar, they still went ahead with the action (Figure 4a). This behavior is in line with the participants forming a commitment to act, which they then carry out. However, the task provided the participants with no reason to abort their action once the opponent had intervened, nor were they instructed to abort their action. Thus, the persistence of movement following an intervention should not be interpreted as a direct test of whether the decoded neural state had already crossed a point-of-no-return for the action. Nevertheless, in the game task, participants not only confirmed their intention to act on 88% of the active brain-based prediction trials (Table 2, Figure 4c,d), but also sometimes accused the opponent of cheating or even claimed agency over the opponent’s actions.

Our results in the game task show that by the time we could decode a person’s intention to act from motor cortex activity, they were almost always (88% of the time) aware of wanting to move. This aligns well with the results of Noel et al. (2025), who showed that the timing of an intention to move co-varied with detectable changes in neural spiking activity across primary motor cortex. Looking at all silent predictions that happened prior to action onset, 12.3% of silent predictions happened early, more than 1 s before action, and 87.7% occurred on time, within 1 s before action (Figure 4c). This distribution closely matched the distribution of intention reports during active prediction trials, where, 12.5% of opponent actions occurred when participants reported no intention to move, and 87.5% occurred when they reported an intention to move (Figure 4d, Table 2). Participants did not know whether a trial would result in an intervention or be silent, so we expect the underlying brain activity up to the prediction to be the same in both trial types. Since the same classifier operated in both silent and active prediction trials, the high correspondence between the two distributions may simply reflect a classifier accuracy of ∼88% after an intervention; counting only predictions that were followed by a positive intention report as correct. Together, these results suggest that interventions happening within the last second prior to action onset were followed by a positive intention report. Similarly, of behavior-based predictions, driven only by a Gaussian policy of reaction times from previous trials, 62.5% were estimated to arrive early, and 37.5% on time. Again, this matched the participants distribution of intention reports, confirming positive intent on 37.1% of these predictions, happening within 1 s of action onset, and absence of intention on the remaining 62.9%, happening more than 1 s in advance of action onset. Both brain-based and behavioral based prediction results support the idea that positive intention is primarily reported within the last second before action.

This still leaves open the source of the early brain-based predictions. One possibility is that they reflect false positives; for example, our decoder may have picked up a mere sub-threshold intention rather than a true neural commitment to act (Schurger et al., 2012). For instance, previous research has shown that participants are inclined to act when the spontaneous neural fluctuations across their motor areas are in a certain phase (Schmidt et al., 2016). Alternatively, early predictions could reflect true pre-conscious neural commitment to act (i.e., one that the participants were still unable to report at the time). In either case, if a brain-based prediction preceded the onset of the subjective awareness of wanting to act, the participants’ intention reports would necessarily be negative. However, intention reports could be biased by the onset of the action itself. Although brain-based predictions preceded the onset of muscle activity on 77.4% of trials, they still happened relatively close to action onset. We cannot rule out that participants based their post-hoc report on whether they actually moved (Verbaarschot et al., 2025). After brain-based interruptions, force exceeded the action threshold in 92.6% of trials with a positive-intent report, but in only 3.6% of trials with a negative-intent report. Perhaps early brain-based predictions happened early enough for participants to (consciously or unconsciously) hold back their action and conclude with a negative intention report.

Interestingly, some participants were not aware of a change in their behavior across tasks. For example, during the scroll task, C2 claimed to be unaware of any changes between behavior-based or brain-based prediction blocks, yet he stopped producing any noticeable wrist force output during brain-based prediction blocks, while maintaining that the images were advancing according to his will. Moreover, the very high percentage of positive intention reports across participants after a brain-based prediction in the game task, and the anecdotal claims of agency over brain-based opponent actions (“I am the robot,” or “I am tipping it [the bucket] on myself”; see Video S5), suggest that there is a window of opportunity in which people take ownership over the resulting actions (Haggard et al., 2002; Moore & Obhi, 2012). In other words, if you get the timing right, actions that appear to be triggered by an opponent in a game can feel like your own.

Our finding that participants overwhelmingly report wanting to move when we decode their intention from motor cortex contradicts an earlier anecdotal report by Grey Walter (Dennett, 1991). In that anecdotal report, participants performed a task similar to our scroll task. Grey Walter purportedly detected participants’ intentions to move via implanted epicortical electrodes in motor cortex and used this to advance a slide projector to the next slide. Participants supposedly reported the unnerving feeling that their mind was being read by the device, as if the projector consistently advanced to the next slide just before they had consciously decided to press the (dummy) button. Grey Walter never formally published these results, likely because they were not properly controlled and lacked ethical approval (Hartmann, 2004), but they nonetheless acquired a mythical status in the volition literature (Dennett, 1991). In our experiment, participants either did not notice the difference between their own action and the classifier-triggered action or appeared to interact effortlessly with the classifier-controlled slideshow (Video S18—19). It is worth noting that our participants were used to controlling both digital and robotic devices through their brain activity in previous experiments spanning multiple years, as they have been part of an ongoing clinical trial that aims to restore autonomy in people with severe paralysis through an implanted BCI. Therefore, they may have been less surprised by actions that were triggered based on their own brain activity than they were the very first time they used a BCI (i.e., the brain may re-calibrate the sense of volition over time). Nevertheless, our results suggest that Grey Walter’s anecdotal results should be viewed with skepticism as we could not replicate his observations.

Participants overwhelmingly lost to the brain-based decoder during the game task. Yet this is not for lack of trying. They all clearly strove to win and reported various strategies intended to undermine being predicted (Video S4, S5). While we observed some changes in neural activity over the course of a session during the game task (Figure 5a-c), the decoder’s accuracy did not deteriorate and participants continued to lose the game throughout the sessions, suggesting that whatever strategies they adopted to act unpredictably to the decoder were insufficient to alter motor cortex activity enough to overcome our real-time system. This supports the interpretation that the detected neural signature was largely unavoidable in the processes leading up to action execution (Schultze-Kraft et al., 2021), consistent with motor cortex being the final cortical stage of neural preparation before action execution (Haggard, 2008). Indeed, any strategy that would successfully undermine our decoder would have had to reduce or otherwise alter activity in the hand-specific region of motor cortex while still enabling execution of the same hand movement—a nontrivial requirement. We therefore think that the changes we saw in neural firing were more likely due to fatigue, disengagement with the task, or the general non-stationarity of the brain, rather than voluntary modulation, because these changes were not linked to action onset and were also visible when the participant was at rest (Figure S9). Previous research has shown that intracortical recordings from primary motor cortex can be unstable, with the statistical properties of the recorded units varying even within a single day (Downey et al., 2018). Shifts in mental state, such as increased frustration or fatigue, can also cause significant changes in neural activity (Myrden & Chau, 2017) and affect BCI performance (Myrden & Chau, 2015), further supporting this interpretation. Despite the above, we did not directly explain to the participants that their brain activity was being used against them and that, to win the game, they must devise a strategy to move without their brain activity being detected by our decoder. However, having all operated a BCI for a long time, we suspected that they might infer it. It would nevertheless be interesting for future studies to test the extent to which such participants could undermine a classifier when explicitly informed how it worked or even using neurofeedback.

Our intention classifier performed similarly well across tasks, in a zero-shot manner, i.e. without additional training, providing evidence for a uniform neural signature of intention. This is despite the clear cognitive differences between a competitive game that was very hard to win and a leisurely scroll through a slideshow at one’s own pace. However, we did observe differences in the neural activity leading up to action onset (Figure 6), which could be due to the nature of the tasks: ranging from purely cue-based (reaction-time task) to completely unrestricted (scroll task) actions (Libet & Gleason, 1982; Verbaarschot et al., 2025; Verbaarschot, Haselager, et al., 2019). That said, we cannot rule out that these differences across tasks may result, at least in part, from general changes in neural firing as a session progressed. In all test sessions, the reaction-time task was used for hardware calibration, before training the classifier and initiating the game and scroll tasks. This introduces an inevitable confound in the experimental design. Without counterbalancing the order of tasks across sessions, we cannot ascertain that these differences are task-related rather than general effects of elapsed time (e.g., due to fatigue). Nevertheless, the differences we observed align well with observations in previous research (Baker et al., 2012; Libet & Gleason, 1982; Salvaris & Haggard, 2014; Travers et al., 2021). In these previous studies, cue-based actions (like our reaction-time task) were preceded by a sharp increase in neural firing, whereas voluntary actions (similar to our game and scroll tasks) were preceded by a more gradual increase in neural firing (Figure 6). Moreover, cue-based actions produced the largest change in neural firing whereas voluntary actions produced the weakest one. Some of these differences may correlate with the exact kinematics of the executed movement (Kaufman et al., 2016). For example, on average, the scroll task showed significantly lower force output and had the lowest change in firing rate prior to action (Figure 6a). Other differences may stem from specific differences in decision making and neural preparation for action between tasks.

Previous attempts at real-time prediction of movement via non-invasive EEG recordings achieved around 60% accuracy, leading to relatively large proportions of false positive and negative predictions (Bai et al., 2011; Han et al., 2020). Similarly, functional Magnetic Resonance Imaging (fMRI)-based decoding strategies achieved marginally significant results (Bode et al., 2011; Soon et al., 2008; Soon et al., 2013). Due to the increased signal-to-noise ratio, intracortical studies have achieved better performances during offline analyses. One study pooled 1019 neurons in premotor cortex areas and anterior cingulate cortex across multiple participants during a self-paced movement task (Fried et al., 2011), revealing a change in neural firing about 500 ms prior to a reported conscious intention to move, which was detectable with 80% accuracy. Another study with individuals with tetraplegia recorded neural activity from posterior parietal cortex, suggesting that motor decisions were encoded 1-0.5 s before awareness (Aflalo et al., 2022). One more study achieved real-time predictions of a binary choice (left/right hand movement) prior to action onset with accuracies up to 68% (Maoz et al., 2012). Here, we demonstrated real-time, brain-based action prediction in an asynchronous setting, achieving a level of accuracy substantially higher (88%) than previously reported. Recent BCI literature has shown an interest in decoding cued motor-preparatory brain activity to make BCIs smoother and easier to control (Rigotti-Thompson et al., 2026). The reliability of our classifier has implications beyond the scientific study of volition, potentially enabling future BCIs to switch between idle and active modes with minimal effort from the user (Bai et al., 2011; Han et al., 2020). This translational potential also raises the question of where along the neural pathway leading up to a voluntary action intention should be decoded.

We set out to ask whether the neural antecedents of voluntary action mark a true neural commitment to act. Indeed, the motor cortical state that we decoded appears to reflect a commitment: it predicted imminent action well, it survived sustained attempts to evade it, and it appeared in the same form whether the action was competitive or performed purely for the participant’s own entertainment. But it arrived only a few hundred milliseconds before movement— and participants could almost always subjectively report wanting to act by the time it did. On this evidence, the interval between decodable commitment and reportable intention is far shorter than the classic picture suggests. However, two interpretations are possible. Either commitment really is settled shortly prior to action onset, and the long antecedents reported over the past six decades reflect some unspecific preparation or a bias toward acting rather than a commitment to act (Schmidt et al., 2016; Schurger et al., 2012). Or action commitment is settled in earlier stages of neural preparation; motor cortex is thought to be the final cortical stage before execution, so a decoder there would encounter the decision of when to act only once it reached the output stage of cortical processing. Teasing these apart requires running the same closed-loop test while recording earlier in the decision process—preempting an action on the basis of parietal or prefrontal activity and asking the actor, in that moment, whether they had yet decided. As BCIs increasingly incorporate regions beyond motor cortex (Aflalo et al., 2022; Gallego et al., 2022; Johnson et al., 2025), this becomes feasible, though it may remain elusive (Schurger & Uithol, 2015). Until then, we can say that at the final cortical stage before movement onset, the decoded intention marks a commitment that the actor already knows about and is followed by movement shortly afterward.

## STAR Methods

## 1. Experimental model and study participants details

This study was part of a multi-site clinical trial, registered at ClinicalTrials.gov (NCT01894802), which aims to demonstrate the safety and efficacy of a bidirectional brain-computer interface that uses implanted intracortical microelectrodes in human participants. Although the data of this study was collected during the participants involvement in the trial, it does not entail any clinical trial outcomes. The study was conducted under an Investigational Device Exemption from the U.S. Food and Drug Administration and ethically approved by the Institutional Review Boards at the University of Pittsburgh and the University of Chicago. Prior to any study procedures, participants provided their informed consent.

Four male participants were tested in this study. Participants were recruited through voluntary research registries for people with spinal cord injury. The sex of participants was based on self-report. As our participant sample included only a single sex, we did not perform any sex-based analyses. We did not assess their gender, as we did not anticipate an effect of gender on our study outcomes. We did not assess the race, ethnicity, or any other socially relevant grouping that our participants may or may not belong to as we did not anticipate an effect of these variables on our study outcomes.

Participant P2 sustained a C5 motor/C6 sensory ASIA B spinal cord injury ten years prior to their participation in the clinical trial and was 25-30 years old when he received the brain implant. P3 sustained a C6 ASIA B spinal cord injury 12 years prior to implantation and was 25-30 years old at the time of implant. C1 sustained a C4 ASIA D spinal cord injury 35 years prior to implantation and was 55-60 years old at the time of implant. C2 sustained a C4 ASIA D spinal cord injury and a right brachial plexopathy 4 years prior to implantation and was 60-65 years old at the time of implant. All participants retained sufficient movement in their right hand to perform a wrist extension.

## 2. Method details*

### 2.1 Data acquisition

Prior to this study, participants were implanted with four 2.4 × 4 mm NeuroPort microelectrode arrays (Blackrock Neurotech, Salt Lake City, UT) in their left hemisphere (Downey et al., 2024; Greenspon et al., 2024). Two 32-electrode arrays were implanted in Area 1 of the contralateral primary somatosensory cortex and two 88-(P2) or 96-(P3, C1, C2) electrode arrays on the precentral gyrus of the motor cortex. Neural data were collected via digital NeuroPlex E headstages using NeuroPort Central software (Blackrock Microsystems, Inc., Salt Lake City, UT). Raw voltages were recorded at 30 kHz, filtered using a 1st-order 750 Hz high-pass Butterworth filter, logged as threshold crossings at -4.5 times the root mean square per electrode, and binned at 50 Hz. Each bin contains the total number of single or multi-unit neural spikes (i.e. threshold crossings) recorded per electrode that happened within that 20 ms window.

Throughout the experiments, participants performed wrist extensions with their right hand. Using this movement, they pushed against a bar that tracked movement performance with a Flexiforce sensor (Tekscan, Inc., Norwood, MA). Force output was recorded via analog input 1 of the main Neural Signal Processor (NSP). To detect movement onset in real-time, a threshold was set for each participant at the start of a session (typically at 5000 a.u.). In addition, movement onset was tracked using an electromyogram (EMG) in at least three sessions per participant. A ground electrode was placed on the wrist bone, and another was placed on the extensor digitorum of the right arm (Figure 2a). In Pittsburgh, EMG data was collected using a Grapevine Nomad (Ripple Neuromed, Salt Lake City, UT) in combination with Trellis software (Pittsburgh). Data was synchronized between the NSP (analog input 3) and Nomad via a sync pulse sent at the onset and offset of a trial, as well as at the registration of movement. In Chicago, the EMG electrodes connected directly to analog input 3 of the NSP via a microelectrode AC amplifier (Model 1800, A-M Systems, Carlsborg, WA). During the reaction-time task, audio output was recorded via analog input 2 on the NSP. All data was formatted and saved for analysis using our own code toolbox in Python 3.7.6 and Matlab 2021a.

### 2.2 Experimental procedures

Participants performed three tasks during the experiment: a reaction-time task, the game task, and a free image scroll task. Each task consisted of a baseline, pre-action, and post-action phase, where actions were marked by prompt right-hand wrist extensions (Figure 1).

#### Reaction-time task

Participants were asked to act as soon as they heard a clearly audible tone (0.2 s, 800 Hz). Each trial started with a 2 s baseline period, during which a black screen was presented to the participant. After that, a white fixation cross appeared, and the audio was presented with a delay drawn from a uniform distribution between 3 and 10 s. After the participant’s action was registered, the fixation cross turned purple and stayed on screen for 0.5 s. Two blocks of 20 trials were presented, with a self-paced break in between.

#### Game task

In this single-player preemption computer game called “Flip-that-Bucket”, participants face a strategic tradeoff, where the potential payoff could increase with their wait time, but if they wait too long, an opponent may preempt them by acting first and seizing the payoff. In this version of the game, participants played a scientist who tried to flip a bucket filling with slime on their robot opponent before the opponent flipped it on them (Figure 1,2, Video S4,S5) (Verbaarschot, Gerrits, et al., 2019). They flipped the bucket by extending their wrist beyond the force threshold. A game consisted of blocks of 20 trials, each culminating in a bucket flip (by the participant or opponent). The player with the most points at the end of the block won the game.

During the game, the opponent tried to predict the participant’s action based on the behavioral timing of their previous actions, or on their ongoing brain activity. After each participant-initiated action, they were asked: “did you want to act?” They could answer with “yes”, “no”, or “maybe”.

For the first five trials of a game, the opponent’s wait time to flip was drawn from a Gaussian distribution with a mean of 8 s and a standard deviation of 2 s. After that, the mean and standard deviation of the Gaussian distribution were set up such that 20% of opponent’s actions would happen 2 s or more before the average scientist move; 60% would take place 2-0 s before the average scientist’s move; and 20% would happen after the scientist moves (Verbaarschot, Gerrits, et al., 2019). Brain predictions, in contrast, were made using a real-time system based on a random forest classifier, which was trained on two blocks of the game (40 trials), where only behavioral predictions were used. For more details on the classification pipeline, see Statistical Analysis below.

Participants completed two blocks (2×20 behavioral trials) of the game task with active behavior-based predictions, followed by validation block (20 behavioral-validation trials). During the validation block, participants played against a behavior-based opponent; but the brain-based classifier was nevertheless making live, though silent, predictions in the background. These silent brain-based predictions enabled us to test the timing and accuracy of our classifier relative to participant action, without interrupting their behavior. In the remainder of the session, we either randomly alternated between active behavior-based and brain-based predictions within a game (3×20 mixed behavioral and brain trials) or played a game with active brain-based predictions only (2×20 brain trials). During an active behavior-based trial in a mixed block, the brain-based prediction was still recorded silently (if it happened before participant or opponent action), and vice versa.

#### Image scroll

In this task, participants used the brain-based classifier from the game task, as is, to scroll through a digital comic book, nature photographs, or Youtube shorts at their own pace (Figure 1, Video S18, S19). Although we always instructed participants to scroll through the digital content by performing a right-hand wrist extension, this was not always used to register and perform the action. In some blocks, we used the force sensor to scroll, whereas in other blocks, we used brain-predictions only. Each test block contained 20-60 trials, where in each trial we presented the participant with a single image or video and waited for their (predicted) response. in behavior-based blocks, a trial concluded when the participant responded. In brain-based blocks, a trial concluded when a brain-based prediction of movement onset was detected by the classifier. During behavior-based blocks, silent brain-based predictions were recorded, and vice-versa. In the final two sessions of C1, we added auditory tones at the time of intention decoding, to enhance awareness of action onset.

#### Data collection

We developed the real-time classification pipeline using short pilot tests with participants P2 (11 sessions) and P3 (6 sessions). During these pilot tests, we focused on collecting training and validation blocks of the game task, using predominantly behavior-based predictions. Once we had established our classification pipeline, we collected the final dataset as reported in this paper, including both behavior-based and brain-based predictions, as well as the reaction-time and scroll tasks. Participant P2 completed 5 sessions of the full experiment, where two sessions included the reaction time task and two sessions included the scroll task. Participant P3 completed 3 sessions of the experiment, two of which included the reaction time task, but none included the scroll task. Participants C1 and C2 completed 3 sessions of the experiment, all including the reaction-time and scroll tasks.

Trials in which participants did not provide their intention report when prompted after an opponent action (game task) or failed to act spontaneously and required a reminder from the experimenter (all tasks) were excluded from analysis. After exclusion, only data from the beginning of the sessions of C2 met our criteria for inclusion. For each session, this data contained 2×20 trials of the reaction time task, 2×20 trials of the game task with behavior-based predictions, 20 trials of validation block of the game task.

### 2.3 Data preprocessing

#### Muscle activity

To estimate the temporal offset between EMG activation and action onset (as measured by the force sensor), we used the reaction-time task. For each participant, all reaction-time trials were aggregated across sessions and time-locked to action onset during offline analysis. At the single-trial level, raw EMG signals were first bandpass filtered between 62 and 250 Hz using a 4th-order Butterworth filter to suppress motion artifacts, baseline drift, and high-frequency noise. The band-passed signal was then mean-centered to remove residual DC offsets and full-wave rectified by taking the absolute value, resulting in a unipolar representation of muscle activation magnitude. To extract a smoother activation envelope, the rectified signal was low-pass filtered using a moving average with a window length of 10 ms, implemented via edge-padded convolution to preserve signal length and reduce boundary artifacts. Finally, the resulting EMG envelope was standardized using per-trial z-score normalization.

EMG onset was estimated as the first time point at which the z-scored EMG signal crossed a threshold of 3 standard deviations within a trial. Then, the temporal difference between this EMG onset and the registered force onset (the latter determined in real-time during the experiment) was computed. The median delay and IQR for each participant were used to report the estimated muscle-to-force latency.

#### Neural multi-unit activity

Neural recordings were processed separately for each participant and session. Single-trials were time-locked to action onset using pre-movement windows of either 1 or 2 s, depending on the analysis (see Figures 3,4 and further methods below). Trials whose duration was shorter than the selected window (1 or 2 s) were excluded from the figures but retained for all statistical analyses. Binned (per 20 ms) spike counts were converted into smooth firing rate estimates by applying a Gaussian filter to each channel independently across time. Filtering was performed with a kernel spanning ±160 ms around each time point (σ = 2 bins), implementing a temporally local weighted average of the signal. At trial boundaries, values required for smoothing beyond the available data were obtained using a symmetric continuation of the signal based on neighboring samples to minimize edge-related artifacts. Smoothed firing rates were then summed across channels, resulting in a one-dimensional Multi-Unit Activity (MUA) profile for each trial. To enable comparisons across participants and recording days, profiles were z-scored within each session using the corresponding mean and standard deviation. All MUA-related figures show the trial-averaged mean, with variability visualized as shaded bands representing ±1 standard deviation, unless otherwise stated.

#### Force activity

One-dimensional force traces were processed for each participant and recording session. During the experiment, force levels above 5000 a.u. would mark the onset of a participant action. For offline analysis, single trials were time-locked to action onset using the same pre-movement windows applied to neural data. To enable comparisons across participants and recording days, we applied trial-wise baseline subtraction using the mean force over the first 500 ms of each trace, and then z-scored the resulting signal within each session using that session’s mean and standard deviation. For visualization, due to the abrupt, non-normal behavior of the signal near action onset, force-related figures show the trial median force, with variability visualized as shaded bands spanning the 25th to 75th percentiles (interquartile range).

## 3. Quantification and statistical analysis

### Real-time brain-based classifier

The real-time decoding framework was designed with the knowledge that preparatory neural activity in the motor cortex plays a clear causal role in the generation of movement intentions (Kornhuber & Deecke, 1985; Shenoy et al., 2003). Accordingly, by decoding neural signatures of the intention to move during movement preparation, the classifier aims to predict when a person commits to action.

At the beginning of each session, a classifier was trained from scratch using neural data collected during the first two blocks of the game task (training blocks), in which participants played against a behavior-based opponent. These blocks provided examples of self-initiated movements, corresponding to trials in which the participant voluntarily flipped the bucket first. The two training blocks provided 25.27 ± 2.69 (mean ± standard deviation) trials per session in which the participant acted first. Given the limited number of training samples available in each session, we selected a random forest classifier due to its ability to model non-linear patterns and mitigate overfitting through ensemble averaging and implicit regularization across trees. This model choice was informed by preliminary analyses conducted on earlier offline sessions. To ensure that inference could be carried out within the real-time constraints imposed by the neural sampling rate (50 Hz), the ensemble size was limited to 200 trees, and the maximum tree depth was capped at 15. The classifier pipeline was made in Python using numpy for batch processing and scikit-learn for model training and real-time inference.

Neural features were derived from sliding windows of spike counts. Binned spike counts (20 ms each) were summed independently for each channel over the preceding 200 ms window (or 10 bins), resulting in a 240-or 256-dimensional (depending on the total amount of implanted microelectrodes per participant) feature vector corresponding to the most recent total population activity at each time step of the trial. These features were computed offline after the two training blocks and served as input to the classifier during training.

Similarly to established supervised learning approaches in intention decoding (Bai et al., 2011; Fried et al., 2011; Lew et al., 2012; Schultze-Kraft et al., 2016; Schultze-Kraft et al., 2021; Schultze-Kraft et al., 2020; Verbaarschot, Gerrits, et al., 2019), we mapped neural activity to intention by labeling the 600 ms interval preceding movement onset as the “pre-movement” (positive) class and an earlier period from −2 s to −600 ms relative to movement onset as the “baseline” (negative) class. This 600 ms window was chosen based on pilot studies that demonstrated that the earliest, average preparatory activity typically departs from baseline approximately within this interval (Figure S3). The −2 s to −600 ms baseline window was selected to ensure a consistent temporal offset to movement onset across trials, given variability in trial duration. Specifically, defining a baseline relative to the beginning of the trial would result in highly heterogeneous temporal offsets from the action (e.g., 2 s in some trials, 4 s or 6 s in others), thereby mixing neural activity at different time periods. By time-locking the baseline window to a later fixed interval preceding movement onset, we ensured that samples were drawn from comparable pre-action periods across all trials. Only these two time windows were used for training; the remaining data of the trial was intentionally excluded to prevent severe class imbalance, which often degrades classifier performance.

Under this labeling scheme, the classifier was effectively trained to answer a binary question: “Will the participant flip the bucket within the next 600 ms?”. After training, the decoder operated continuously and asynchronously in the background. For each 20 ms bin, the features were computed from the most recent 200 ms buffer of neural activity, and the classifier generated a continuous confidence score corresponding to the fraction of trees predicting an imminent action (i.e., action within the next 600 ms). The score was tested against the default 0.5 threshold to determine the binary classification output (movement within the next 600 ms or not). This procedure enabled real-time monitoring of movement intention based solely on the ongoing neural stream.

### Estimating electrode importance for decoding

To identify the most informative electrodes for decoding, a random forest classifier was trained offline using the identical parameter configuration to the real-time experiments. Training was performed using the first two blocks of the game task (training blocks) and repeated 500 times; each initialized with a different random seed. At each iteration, the random forest’s feature importance values (one per electrode) were extracted from the trained model. These values quantify how strongly each electrode contributed to reducing classification error and improving predictive performance. Feature-importance scores were then averaged across iterations, obtaining a robust estimate of each electrode’s salience. This procedure was carried out separately for each session and participant.

Electrode importance was subsequently related to spatial location on the NeuroPort microelectrode arrays by averaging importance values across sessions within each participant. This resulted in a single importance-score per electrode and participant, reflecting its overall contribution to decoding performance and enabling comparisons across anatomical locations.

### Opponent performance evaluation

Opponent performance was evaluated separately for the brain-based classifier and the behavior-based predictions using two modes of operation: (1) silent predictions, in which the model made a prediction in real-time, but this prediction did not affect gameplay (i.e. a prediction that the participant is about to move did not trigger the opponent to flip the bucket); and (2) active predictions, in which the opponent intervened during gameplay by flipping the bucket when it calculated that participant was about to act.

For the brain-based classifier, silent predictions were quantified using the continuous model output logged in the background during gameplay. Specifically, for each trial in which the participant performed an action without being interrupted, we extracted the sequence of silent brain-based predictions spanning the trial. To mimic the real-time behavior of the classifier, the first positive-class output within the trial was defined as the time at which the opponent would have flipped the bucket had the classifier been engaged. We then computed the time difference between this silent brain-based prediction onset and the participant’s action onset, indicating how far in advance the decoder predicted the participant’s action.

In line with the original Flip-that-Bucket electroencephalography (EEG) experiment (Verbaarschot, Gerrits, et al., 2019), each trial was categorized into one of three categories. Predictions occurring earlier than 1 s before action onset were classified as “early.” Predictions occurring within the 1 s interval leading up to action onset were classified as “on-time.” And predictions occurring after action onset were classified as “late.” For each category, the percentage of trials in that category was computed as well as the mean and standard deviation of prediction timing.

In contrast to the brain-based classifier, the behavior-based opponent did not log a continuous background prediction stream and therefore does not permit an empirical “silent prediction” timing analysis. However, because the behavior-based opponent operates according to a Gaussian timing policy derived from participants’ wait times, the distribution of its action timings across the same early/on-time/late categories can be estimated from the way the distribution was devised (see Experimental Procedures). Hence, −2 s corresponds to the 20th percentile of the Gaussian distribution, and 0 s corresponds to the 80th percentile. Because a Gaussian distribution is symmetric, the median, which is also the mean, lies halfway between these two percentiles, resulting in a mean action time of −1 s relative to action onset. Using our category definitions, −1 s corresponds to the mean of this distribution, thus 50% of actions are expected to occur earlier than −1 s (“early”). By construction, 20% of actions are expected to occur after action onset (“late”). The remaining probability mass between −1 s and 0 s therefore corresponds to 30% of actions (“on-time”). This 50-30-20% split represents the expected distribution of the behavior-based model across the three interval-categories. The empirical timing-category frequencies produced by this type of opponent are therefore expected to approximate these proportions (up to sampling and goodness-of-fit variability).

Because active predictions interrupted the participant during gameplay (i.e., the opponent flipped the bucket first, meaning that the participant typically did not move), it was not possible to measure the prediction onset time relative to the participant’s action onset on those trials. Instead, active predictions—both brain- and behavioral-based—were evaluated using participant’s response to the intention question presented immediately after each opponent intervention. Participants were asked “Did you want to act?” and could provide one of three responses: “yes,” “no,” or “maybe.” “Maybe” responses accounted for 0.5% of trials on average and were therefore excluded from analysis due to their rarity.

In brain-based prediction blocks (where the participants played exclusively against the brain-based classifier), we computed the classifier win rate as the percentage of trials in which the classifier flipped the bucket before the participant acted. This measure indexes classifier responsiveness and was later compared to the proportion of conceded interventions (i.e., participants confirming they wanted to act).

### Comparing neural and force activity preceding opponent actions

Opponent actions were related to participants’ reported intention by analyzing neural and force activity preceding opponent-initiated bucket flips (active predictions) during the game task. Within each session, trials were stratified by intention report (“yes” vs “no”) and opponent type (brain-based vs behavior-based). Activity was time-locked to the opponent’s action (bucket flip), and 2 s of pre-action neural and force data were extracted. The signal-specific preprocessing steps described in section “2.3 Data processing” were applied to each recording modality. Summary statistics were then computed separately for each combination of opponent type and intention report (brain-based/yes, brain-based/no, behavior-based/yes, behavior-based/no).

For the statistical comparison of neural activity between brain- and behavior-based predictions, we included trials with at least 320 ms of neural data preceding the opponent’s action. For each trial, mean z-scored MUA was calculated over the final 200 ms before the action, matching the neural activity window used by the decoder for classification. Brain- and behavior-based trials were compared separately for each participant and intention response (“yes” or “no”) using two-sided Welch’s t-tests. P-values were Bonferroni-corrected across the six comparisons (three participants × two intention responses).

### Correlating silent prediction timing to intention reports

To relate silent prediction timing to intention reports from active predictions, we first note that “late” silent predictions would not have triggered an intention question in the actual task, because the participant would have acted first and beaten the opponent. Therefore, only “early” and “on-time” silent predictions were considered relevant for comparison with intention reports.

For each participant, the “early” and “on-time” silent proportions were normalized by their sum, such that on-time / (on-time + early) represents the estimated proportion of potential interventions that would have occurred within 1 s of action onset, and early / (on-time + early) represents the estimated proportion of potential interventions that would have occurred earlier than 1 s before action onset. These normalized values were computed at the participant level and aggregated across participants. The proportions were then compared to the distribution of intention reports (“yes” vs. “no”) obtained during active predictions. Although silent and active predictions were derived from different trials, this comparison assumed that both prediction types followed a similar timing relative to participant action, since they were generated by the same classifier. The analysis provided a principled way to relate background decoding timing and accuracy to participants’ explicit intention reports.

### Regression-based characterization of neural dynamics across time

Trial-by-trial changes in brain activity over the duration of the session for the game task were quantified using a regression analysis, performed separately for each participant and session. The analysis used time-locked pre-movement activity from all participant-initiated actions, excluding trials with insufficient pre-movement duration (< 1s).

Spiking activity was transformed into smooth firing rate estimates by applying a Gaussian filter (STD = 2) across time to each channel independently. Binned and filtered spike counts were converted to firing rate by dividing them by the bin width (20 ms). Rare, high-amplitude transients potentially harmful to regression estimates were mitigated by excluding anomalous trials using a robust z-scoring procedure for outlier detection (Rousseeuw & Hubert, 2011). For each trial, firing-rate activity was first averaged across channels, bin by bin, to obtain a single-trial temporal profile. At each time bin, activity was normalized across trials using the median and the median absolute deviation (MAD; scaled by 1.4826 to approximate a standard-deviation-like metric). Trials whose absolute robust z-score exceeded 3 at any time bin were excluded from subsequent analyses.

Within-session dynamics were assessed by sorting trials according to their occurrence (i.e., the first trial in the session, the second trial in the session, and so on across blocks) and indexing them by trial number (Figure 5a). We fit a linear regression model, across all the trials in a session, to each channel and each time bin within the pre-movement window. We did this to predict firing rate as a function of trial index. The slope of the model captured systematic linear changes in activity within the session, producing a two-dimensional slope map (channels × time) summarizing within-session temporal trends (Figure 5a).

Potentially meaningful slopes were separated from noise by estimating a session-specific null scale using the MAD-based robust z-scoring method (Rousseeuw & Hubert, 2011), computed session-wise across all channels and time bins. Slopes with |z| > 2 were treated as having substantial magnitude. Then, bins with z>2 or z<-2 were encoded as “strong positive evidence” or “strong negative evidence,” respectively, while the remaining bins were discarded from further analysis. A binomial statistical test (p = 0.5, alpha = 0.05) was then applied independently to each channel over the surviving time bins to determine whether positive evidence significantly exceeded negative evidence (“predominantly positive”) or vice versa (“predominantly negative”). This test was used as a directional classification heuristic rather than a strict inferential procedure. Channels without significant positive or negative dominance were labeled “null,” reflecting activity lacking a reliable direction. This pipeline produced three electrode clusters based on their slope sign consistency (Figure 5b,c).

Cluster-level dynamics were visualized by summarizing within-session activity separately for the positive-and negative-slope clusters. Specifically, the average firing rate of each channel group was computed on single trials, then z-scored within each session using the session-specific mean and standard deviation. Single-session trials were plotted from earliest to latest using a gradient palette to indicate trial order, thereby illustrating how firing rates evolved over the session (Figure 5b,c). To determine whether the channel-group patterns identified from pre-movement activity also extended to another task-irrelevant epoch, we applied the same electrode clusters to firing rates during a separate 1-second baseline period time-locked to trial onset. This allowed us to assess whether the channels classified according to their pre-movement within-session trends showed similar directional activity patterns during a period of time unrelated to action preparation.

Finally, the relationship between cluster-level neural shifts and intention decoding was assessed using feature importance estimates from the random forest classifier. Because importance values are normalized to sum to unity, they quantify each electrode’s fractional contribution to predictive performance. Importance values were mapped and aggregated by cluster to compute total cluster importance, interpreted as the functional weight of the group. In addition, cluster size was quantified as the proportion of electrodes in each cluster relative to the total recorded electrodes, enabling direct comparison between a cluster’s decoding contribution and its relative size.

### Task-dependent differences in pre-movement neural activity

To compare differences in neural activity prior to participant action during the reaction-time, game and scroll tasks, we performed two complementary analyses based on single-trial summaries of the pre-movement MUA time course. These analyses independently quantified differences in response magnitude and temporal profile, respectively, and were applied to all available pairwise task contrasts for each participant (P2 and C1: reaction vs. game, reaction vs. scroll, and game vs. scroll; P3: reaction vs. game).

First, to quantify differences in response magnitude, each trial was summarized by its peak pre-movement activity, defined as the 95th percentile of the z-scored MUA within the 1-second window preceding movement onset. Rather than using the absolute maximum, we used this definition to better capture the upper range of the pre-movement response while reducing sensitivity to brief, potentially artifact-driven fluctuations.

Second, to quantify differences in the temporal profile of pre-movement activity independent of overall magnitude, we used the area under the curve (AUC) of the normalized pre-movement MUA trace as a shape-based summary measure. For each trial, the pre-movement MUA trace was min-max normalized to the range of 0 to 1, and the AUC was computed using the trapezoidal rule (numerical integration) across time bins. Under this normalization, higher AUC values indicate that activity rose earlier and remained elevated over more of the pre-movement window, whereas lower AUC values indicate that activity stayed low for most of the window and increased only closer to movement onset.

For both the peak-MUA and AUC analyses, overall task effects within each participant were first assessed using Kruskal-Wallis tests. When the omnibus test was significant, post hoc pairwise task comparisons were performed using two-sided Mann-Whitney tests, with Bonferroni correction for multiple comparisons. Corrected post hoc p-values were reported using standard significance codes (* p < 0.05, ** p < 0.01, *** p < 0.001, **** p < 0.0001; otherwise non-significant, ns) and visualized as bracket annotations above boxplots of trial-wise peak MUA and AUC values for each participant.

### Identifying slow- and fast-rise onsets of neural activity

To identify distinct stages in the pre-movement ramp up of neural activity, we applied a breakpoint analysis using segmented regressions to the trial-averaged, z-scored MUA profile of each task. Before running the analysis, each MUA profile was min-max normalized to the range of 0 to 1 (as seen in Figure 6a, middle figure). This normalization removed differences in overall response magnitude while preserving the temporal shape of the activity profile, allowing onset estimates to be determined independently of scale.

Each normalized profile was fitted with a single continuous piecewise-linear model composed of three connected regression segments. The first segment was a constant regression with an estimated intercept and a slope fixed at zero, representing the initial baseline level of neural activity. The second segment was a linear regression with a positive slope, representing a gradual or slow increase in activity. The third segment was a second linear regression with a steeper positive slope, representing the subsequent fast increase in activity. The model was fit as a single constrained function, such that all three segments were estimated simultaneously and forced to join at two breakpoint locations, ensuring continuity of the fitted activity without any discontinuities at the transitions:

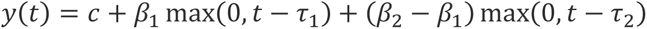

Where *c* is the baseline level, *τ*_1_and *τ*_2_ are the onset times of the slow and fast rising phases, respectively, and *β*_1_ and *β*_2_ are their corresponding slopes. The coefficient on the second term, (*β*_2_ −*β*_1_), represents the change in slope at *τ*_2_, ensuring that *y*(*t*) increases at rate *β*_1_ between *τ*_1_and *τ*_2_, and at a rate *β*_2_ thereafter while remaining continuous at both breakpoints.

The first breakpoint, marking the transition from the constant baseline segment to the slow positive slope, was defined as the “slow-rise onset”. The second breakpoint, marking the transition from the slow-rise segment to the steeper positive slope, was defined as the “fast-rise onset”. To ensure that the model represented the expected progression from baseline to slow and then fast activity buildup, the slopes of both rising segments were constrained to be positive, and the slope of the final segment was required to be greater than that of the preceding segment. Each of the three segments was also required to span at least 100 ms, corresponding to five 20-ms time bins.

The segmented model was fitted from 1 s before movement onset until the peak of the normalized, trial-averaged MUA profile, with the peak identified as the maximum value within the period beginning 300 ms before movement. Ending the fitting window at the peak prevented the subsequent decline in activity from influencing the estimated onsets. Candidate pairs of slow- and fast-rise breakpoints were evaluated exhaustively at 20-ms intervals across the fitting window. For a complete 1-s pre-movement window containing 50 bins, this procedure provided an average of 41 possible breakpoint locations and 807 temporally ordered pairs of breakpoint locations, of which 651 satisfied the requirement that all three segments span at least five bins. Because the fitting window ended at the task-specific peak, the exact number of breakpoint locations and candidate pairs could vary across tasks. Candidate pairs that did not produce a positive slow-rise slope followed by a steeper positive fast-rise slope were excluded.

For each remaining candidate pair, the intercept and slopes of the three connected segments were estimated jointly. The final breakpoint pair was selected as the valid solution that minimized the sum of squared residuals between the observed MUA profile and the complete three-segment fit. Thus, the resulting slow- and fast-rise onsets represented the globally best-fitting breakpoint locations on the temporal bin grid.

## Acknowledgements

We thank the clinical trial participants for their continuous and enthusiastic participation in our research. Without them, none of this would be possible. In addition, we would like to thank Tyler Simpson for creating the wrist-extension measurement device, and Jeffrey Weiss for his never-ending patience and technical support on the development of the real-time decoding pipeline. We would also like to thank Carleigh May, Debbie Harrington and Ashley van Driesche for their well-organized support in this multi-site clinical trial. Finally, we would like to thank Pim Haselager, Jason Farquhar and Anne Gerrits for their knowledge, enthusiasm and continued support that brought Flip-that-Bucket into existence.

This research was supported by the Dutch Research Council (NWO Rubicon: 019.193SG.011) and the National Institute of Neurological Disorders and Stroke of the National Institutes of Health (NIH) and the NIH BRAIN Initiative under Award Number UH3NS107714. The content is solely the responsibility of the authors and does not necessarily represent the official views of the National Institutes of Health.

## Author contributions

C.V., A.M., and U.M. contributed to the experiment design. C.V. and A.M. implemented the experiment. C.V. and A.A. performed data acquisition. C.V. and A.M. performed data analysis under the supervision of U.M., J.C., and R.G. C.V and A.M. wrote the paper draft and C.V, A.M., A.A., U.M., J.C. and R.G. contributed to the paper revisions.

## Declaration of interests

R.G. is on the scientific advisory board of Neurowired LLC and had a consulting role with Blackrock Neurotech during this project. R.G. and J.L.C. received research funding from Blackrock Neurotech, Inc. though that funding did not support this project. C.G. is a consultant for Neuralink. U.M. serves as a consultant to SynthBee, Inc. on the development and evaluation of intent architectures in artificial systems. The remaining authors declare no competing interests.

## Declaration of generative AI and AI-assisted technologies

No generative AI or AI-assisted technologies were used in preparation of this scientific publication.

## Data availability

Upon reasonable request to the study PI, the de-identified data that support the findings of this study can be made available. Data access can be requested from the Data Archive BRAIN Initiative (DABI) under project code VB7674Y4UPDW at http://doi.org/10.18120/zryz-cs73.

## Code availability

Customized code used for analysis is available through GitHub at github.com/CeciVerbaarschot/Preempting_intended_movement.git. The implementation of the Flip-that-bucket game can be found at github.com/jadref/buffer_bci/tree/master/matlab/movementBCIgame/FlipThatBucket.

## Notes

http://doi.org/10.18120/zryz-cs73

https://github.com/CeciVerbaarschot/Preempting_intended_movement

https://github.com/jadref/buffer_bci/tree/master/matlab/movementBCIgame/FlipThatBucket

