## Supplementary material for "Real-time intracortical decoding from human motor cortex reveals a task-independent commitment to act that aligns with conscious intent": Figure S1

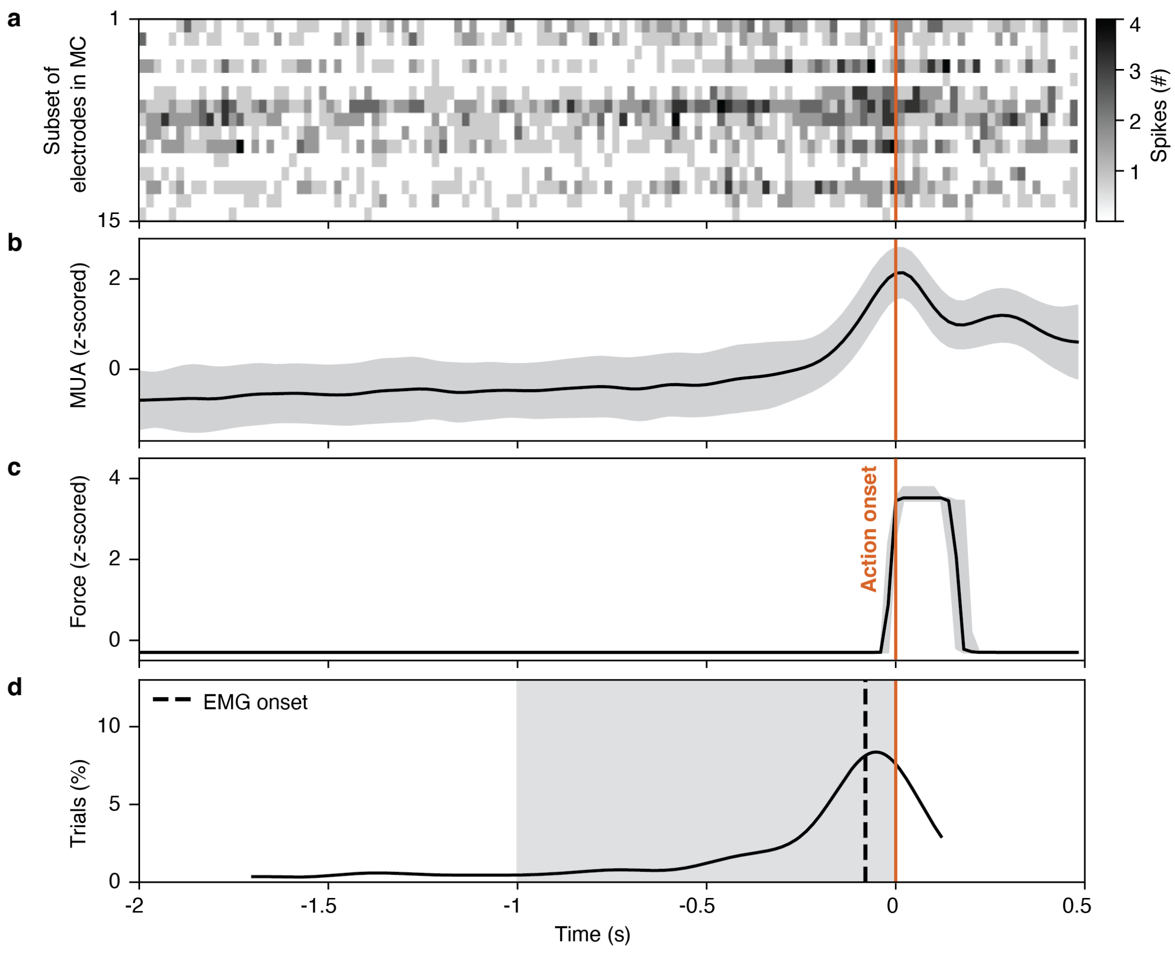


***Figure S1.*** *Neural preparation for action and classifier performance during the game task with participant C2. (a) A typical raster plot, showing the number of recorded threshold crossings in every 20ms time bin 2000ms before to 500ms after action onset (time 0) across a subset of 15 recorded microelectrodes. (b) Mean and standard deviation of the smoothed firing rates across all electrodes. (c) Median and 25-75 percentile ranges of the force output recorded around action onset (time 0). (d) Percentage of silent brain-based classifier predictions happening around action onset (time 0). The time range of on-time predictions is marked in grey. The median onset of muscle activity (EMG onset) based on the reaction-time task is reported.*
