## Supplementary material for "Real-time intracortical decoding from human motor cortex reveals a task-independent commitment to act that aligns with conscious intent": Figure S2

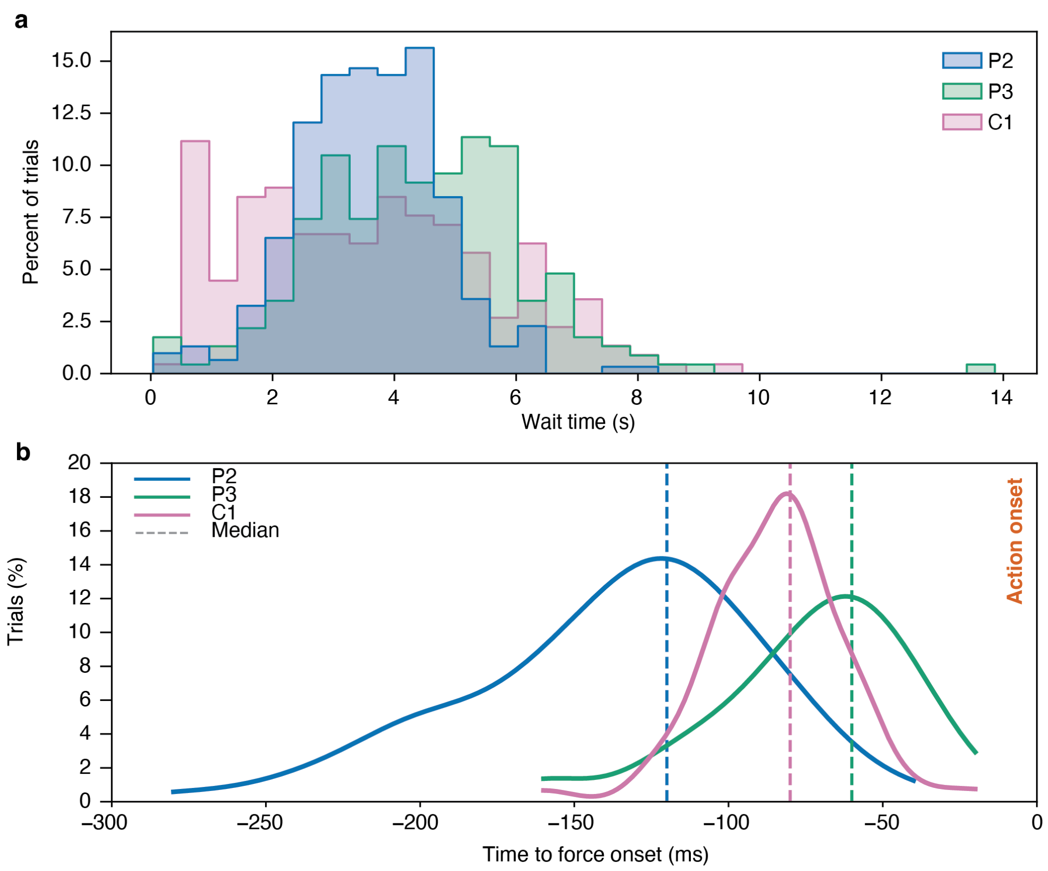


***Figure S2.*** *Overview of the participants' wait times and muscle onset relative to action. (a) Overview of wait times during the game task. Wait time indicates how long (in s) each participant waited before acting in the game task. (b) Overview of onset of muscle activity relative to action, as measured by a considerable increase in force (time 0) during the reaction-time task. The median onset of muscle activity per participant relative to action is indicated with a dotted vertical line.*
