## Supplementary material for "Real-time intracortical decoding from human motor cortex reveals a task-independent commitment to act that aligns with conscious intent": Figure S3

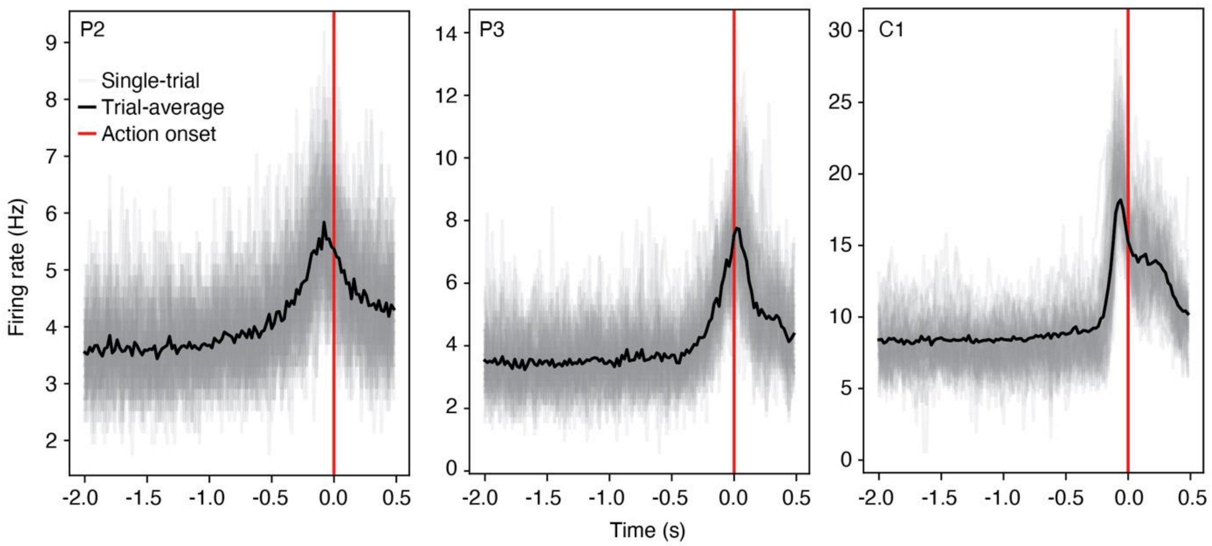


***Figure S3.*** *Single-trial neural responses preceding action onset (time 0) during the game task. Gray traces show the average raw firing rate across electrodes for individual trials, computed in 20 ms bins; the black trace shows the trial-averaged response. Across participants, neural activity exhibits a clear increase beginning approximately 500 ms before action onset.*
