## Supplementary material for "Real-time intracortical decoding from human motor cortex reveals a task-independent commitment to act that aligns with conscious intent": Figure S6-7

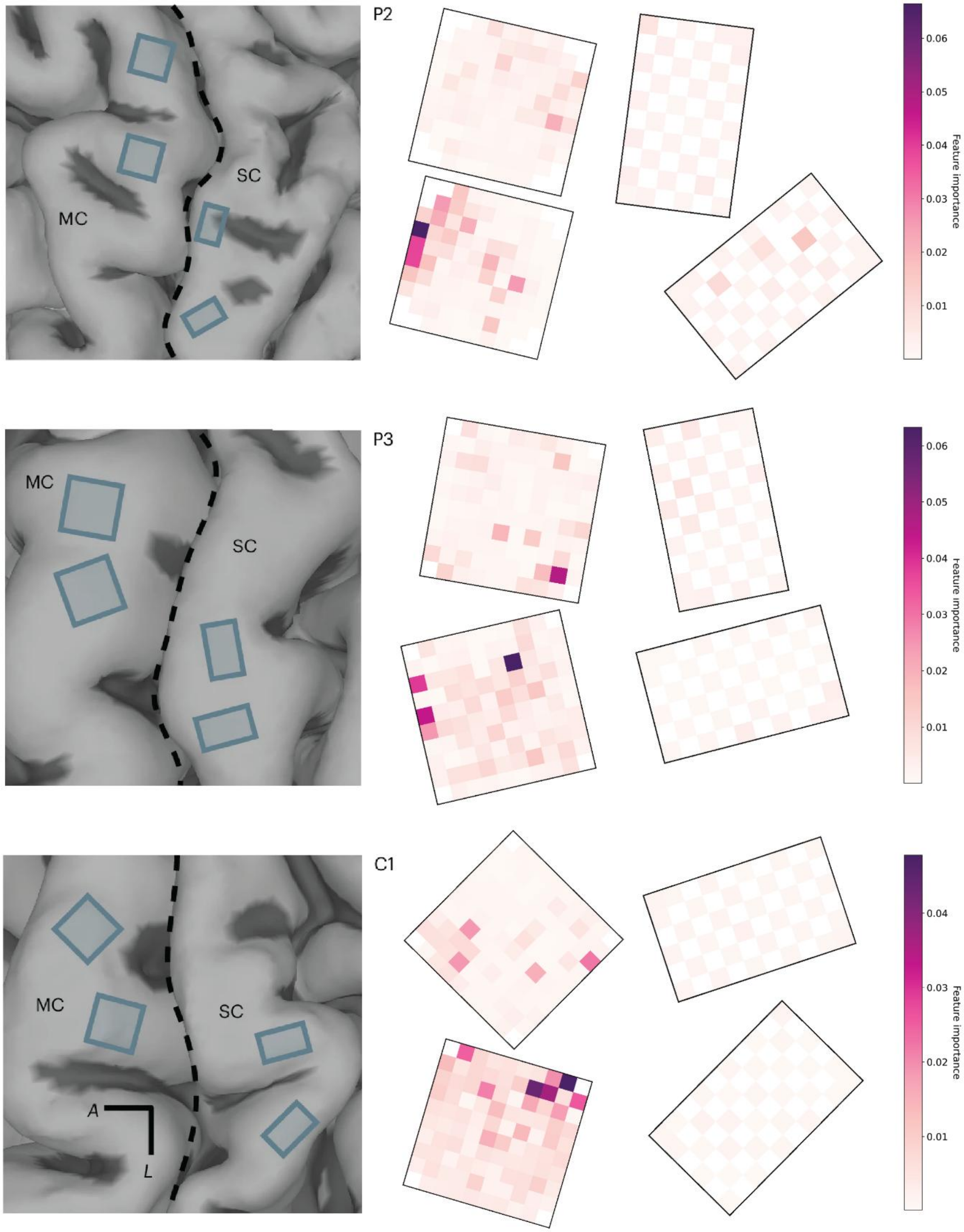


***Figure S6.*** *Electrode importance for the random-forest classifier that was trained to detect the participant's intention to act during the game task. The left column shows the implant locations of the microelectrode arrays in the motor cortex (MC) and somatosensory cortex (SC) (images adapted from Greenspon et al, 2025). For reference, the central sulcus is highlighted with a dotted line. The right column shows how much each microelectrode contributed to*

*classifier performance, on average (feature importance; computed across sessions). For all three participants, a select cluster of electrodes on the motor cortex arrays dominate the relative feature importance of the trained classifier.*

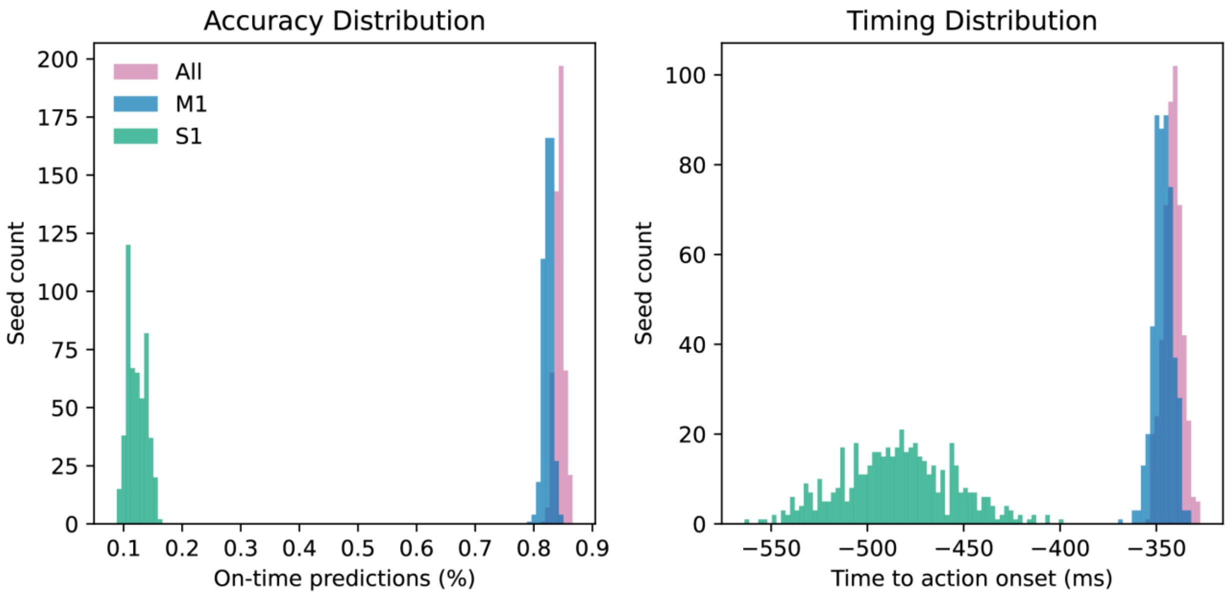


***Figure S7.*** *Offline asynchronous intention prediction using electrode subsets. A random forest classifier was trained offline using the game data acquired during the real-time sessions while using the same hyperparameters as the online decoder. For each participant, the model was trained on the training blocks and evaluated on uninterrupted participant-initiated actions from the test blocks. An asynchronous prediction was considered successful if it occurred within 1 s prior to movement onset. For each successful prediction, the time from action onset was recorded. For each electrode configuration (All, M1, and S1), the classifier was independently trained and tested 500 times using different random seeds. The same set of 500 random seeds was used across all three electrode configurations to ensure that differences reflected electrode selection rather than stochastic variation in model training. Histograms show the distributions of the percentage of on-time predictions (left) and mean prediction time relative to action onset (right) across the 500 model initializations for each electrode configuration. Models trained using all electrodes achieved the highest performance (84.40 ± 0.79% on-time predictions, -341.55 ± 4.89 ms; mean ± standard deviation), followed closely by models trained using M1 electrodes alone (82.42 ± 0.78%, -346.54 ± 4.89 ms). In contrast, models trained using only S1 electrodes achieved substantially lower performance (12.25 ± 1.65%, -484.55 ± 28.94 ms), suggesting that predictive information was concentrated primarily in motor cortex.*
