## Supplementary material for "Real-time intracortical decoding from human motor cortex reveals a task-independent commitment to act that aligns with conscious intent": Figure S9

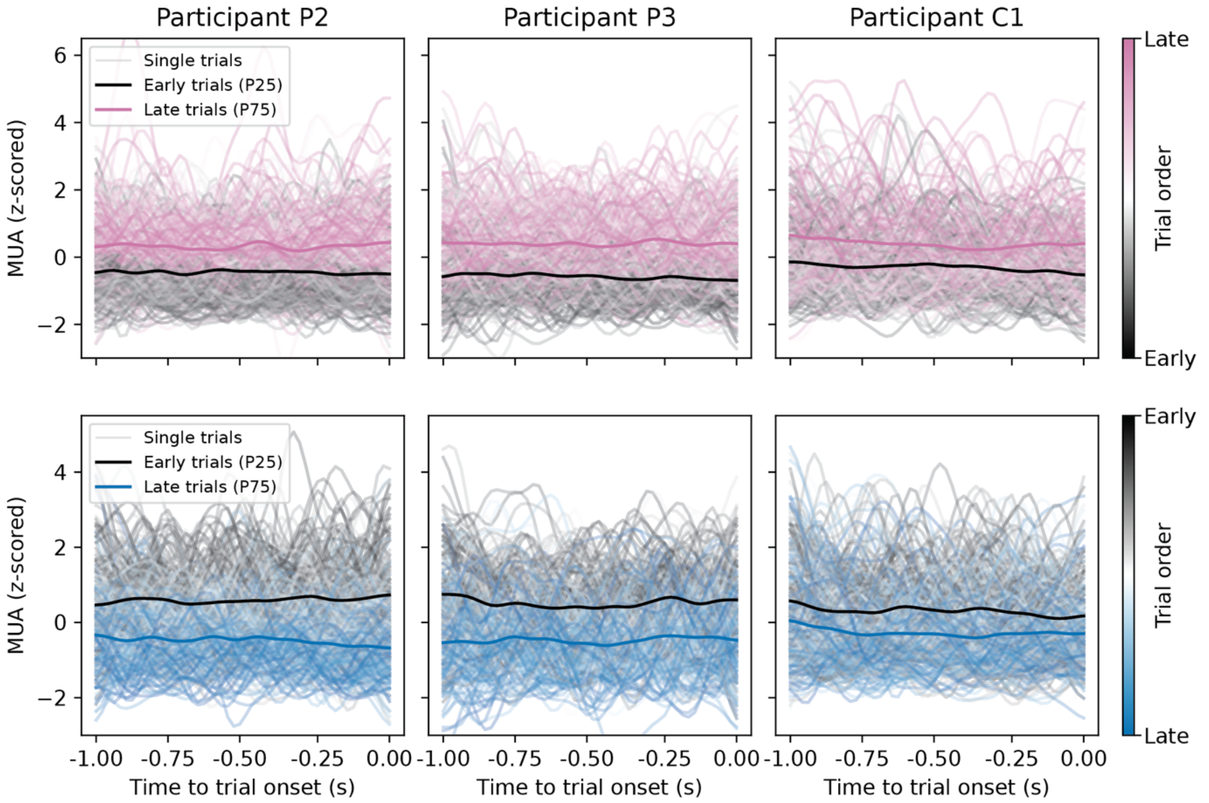


***Figure S9.*** *Baseline activity in the final second before trial onset reflects the same cluster-specific within-session patterns observed in pre-movement dynamics. Electrode clusters were defined based on regression slopes of pre-movement activity (see Methods: Regression-based characterization of neural dynamics across time) and then applied to firing rates during a separate 1 s baseline period time-locked to trial onset (time 0). For each cluster, firing rates were averaged across electrodes on single trials and z-scored within session. Thin lines show single-trial activity, colored by trial order (early to late within session), and thick lines show the trial-averaged response for early (25th percentile) and late (75th percentile) trials. Across participants, channels classified as exhibiting positive within-session trends during pre-movement activity (top row) also show increasing baseline activity over the session, and channels with negative trends (bottom row) show decreasing baseline activity. These results indicate that the cluster-specific modulation identified during action preparation generalizes to a task-unrelated baseline period.*
