## Supplementary material for "Real-time intracortical decoding from human motor cortex reveals a task-independent commitment to act that aligns with conscious intent": Table S8

***Table S8.*** Welch's t-test results comparing mean z-scored MUA during the 200 ms preceding opponent action onset between brain- and behavior-based predictions, separately for "yes" and "no" intention reports. Significance codes: ns, p ≥ .05; *p < .05; **p < .01; ***p < .001; ****p < .0001.

| Participant | Intention | Brain-based (mean) | Behavior-based (mean) | Difference | Statistic (t) | DF | P-value (raw) | P-value (adjusted) | Significance |
|---|---|---|---|---|---|---|---|---|---|
| P2 | Yes | 1.193 | 0.453 | 0.74 | 4.22 | 29.4 | 2.18E-04 | 1.00E-03 | ** |
| P2 | No | 0.446 | -0.339 | 0.785 | 3.25 | 19.2 | 4.00E-03 | 2.50E-02 | * |
| P3 | Yes | 0.535 | 0.123 | 0.412 | 3.13 | 48.6 | 3.00E-03 | 1.80E-02 | * |
| P3 | No | 0.204 | -0.286 | 0.49 | 4.65 | 28.2 | 7.11E-05 | 4.27E-04 | *** |
| C1 | Yes | 0.641 | 0.188 | 0.453 | 3.65 | 67.5 | 5.11E-04 | 3.00E-03 | ** |
| C1 | No | 0.454 | -0.289 | 0.743 | 3.49 | 63 | 8.93E-04 | 5.00E-03 | ** |
