## Supplementary material for "Real-time intracortical decoding from human motor cortex reveals a task-independent commitment to act that aligns with conscious intent": Table S10

***Table S10.*** *Size of slope-based electrode clusters and their relative contribution to the random forest's feature importance scores.*

| | Comparison | P2 | P3 | C1 | Average |
|---|---|---|---|---|---|
| Positive | Importance | 18.95% | 20.62% | 20.98% | 20.2% |
| | Electrodes | 11.72% | 16.41% | 14.06% | 14.1% |
| Negative | Importance | 43.88% | 51.47% | 49.93% | 48.4% |
| | Electrodes | 23.96% | 19.14% | 25.26% | 22.8% |
| Null | Importance | 37.17% | 27.92% | 29.10% | 31.4% |
| | Electrodes | 64.32% | 64.45% | 60.68% | 63.2% |
