## Supplementary material for "Real-time intracortical decoding from human motor cortex reveals a task-independent commitment to act that aligns with conscious intent": Table S11

***Table S11.*** *OLS linear regression results assessing changes in silent on-time prediction percentage and prediction timing across chronological blocks, including participant-adjusted overall trends, participant-specific slopes, and block × participant interaction tests. Significance codes: ns, p ≥ .05; *p < .05; **p < .01; ***p < .001; ****p < .0001.*

| Outcome | Analysis | Pp. | Model | Test | Slope | Units | Stat. | Statistic Value | DF | DF Den | P-value | R Squared | Adjusted R Squared | Sign. |
|---|---|---|---|---|---|---|---|---|---|---|---|---|---|---|
| On-time percentage | Participant-adjusted common slope | All | outcome ~ block + C(participant) | OLS slope t-test | 1.46 | percentage points/block | t | 3.17 | 39 | - | 2.94E-03 | 0.27 | 0.22 | ** |
| On-time percentage | Participant-specific slope | P2 | outcome ~ block | OLS slope t-test | 0.89 | percentage points/block | t | 1.32 | 11 | - | 2.14E-01 | 0.14 | 0.06 | ns |
| On-time percentage | Participant-specific slope | P3 | outcome ~ block | OLS slope t-test | 2.00 | percentage points/block | t | 2.14 | 13 | - | 5.19E-02 | 0.26 | 0.20 | ns |
| On-time percentage | Participant-specific slope | C1 | outcome ~ block | OLS slope t-test | 1.30 | percentage points/block | t | 1.79 | 13 | - | 9.64E-02 | 0.20 | 0.14 | ns |
| On-time percentage | Block x participant interaction | All | outcome ~ block + C(participant) vs. outcome ~ block * C(participant) | Nested-model F-test | - | - | F | 0.46 | 2 | 37 | 6.36E-01 | 0.29 | 0.20 | ns |
| Timing | Participant-adjusted common slope | All | outcome ~ block + C(participant) | OLS slope t-test | -18.50 | ms/block | t | -3.97 | 39 | - | 2.98E-04 | 0.29 | 0.24 | *** |
| Timing | Participant-specific slope | P2 | outcome ~ block | OLS slope t-test | -27.25 | ms/block | t | -3.81 | 11 | - | 2.91E-03 | 0.57 | 0.53 | ** |
| Timing | Participant-specific slope | P3 | outcome ~ block | OLS slope t-test | 3.92 | ms/block | t | 0.64 | 13 | - | 5.31E-01 | 0.03 | -0.04 | ns |

| | | | | | | | | | | | | | | |
|---|---|---|---|---|---|---|---|---|---|---|---|---|---|---|
| Timing | Participant-specific slope | C1 | outcome ~ block | OLS slope t-test | -35.24 | ms/block | t | -5.36 | 13 | - | 1.31E-04 | 0.69 | 0.66 | *** |
| Timing | Block x participant interaction | All | outcome ~ block + C(participant) vs. outcome ~ block * C(participant) | Nested-model F-test | - | - | F | 10.92 | 2 | 37 | 1.87E-04 | 0.55 | 0.49 | *** |
