## Supplementary material for "Real-time intracortical decoding from human motor cortex reveals a task-independent commitment to act that aligns with conscious intent": Table S12-17

***Table S12.*** *Kruskal–Wallis test results comparing differences in pre-movement peak MUA between experimental tasks for each participant.*

| Participant | Comparison | DF | Statistic (H) | P-value | Significance |
|---|---|---|---|---|---|
| P2 | Reaction vs Game vs Scroll | 2 | 149.32 | 3.76E-33 | **** |
| P3 | Reaction vs Game | 1 | 15.09 | 1.02E-04 | *** |
| C1 | Reaction vs Game vs Scroll | 2 | 164.32 | 2.08E-36 | **** |

***Table S13.*** *Post hoc pairwise Mann–Whitney test results comparing differences in pre-movement peak MUA between experimental tasks for each participant. P-value is adjusted using Bonferroni-correction.*

| Participant | Comparison | Statistic (U) | P-value (raw) | P-value (adjusted) | Significance |
|---|---|---|---|---|---|
| P2 | Game vs Scroll | 7889 | 6.23E-06 | 1.87E-05 | **** |
| P2 | Reaction vs Game | 21829 | 2.08E-29 | 6.23E-29 | **** |
| P2 | Reaction vs Scroll | 2805 | 3.68E-16 | 1.10E-15 | **** |
| P3 | Reaction vs Game | 6208 | 1.03E-04 | - | *** |
| C1 | Game vs Scroll | 18685 | 8.02E-31 | 2.41E-30 | **** |
| C1 | Reaction vs Game | 9553.5 | 5.63E-03 | 1.69E-02 | * |
| C1 | Reaction vs Scroll | 7997 | 6.82E-26 | 2.05E-25 | **** |

***Table S14.*** *Kruskal–Wallis test results comparing differences in the pre-movement area under the curve of neural profiles between experimental tasks for each participant.*

| Participant | Comparison | DF | Statistic (H) | P-value | Significance |
|---|---|---|---|---|---|
| P2 | Reaction vs Game vs Scroll | 2 | 121.47 | 4.19E-27 | **** |
| P3 | Reaction vs Game | 1 | 22.76 | 1.84E-06 | **** |
| C1 | Reaction vs Game vs Sroll | 2 | 88.05 | 7.59E-20 | **** |

***Table S15.*** *Post hoc pairwise Mann–Whitney test results comparing differences in the pre-movement area under the curve of neural profiles between experimental tasks for each participant. P-value is adjusted using Bonferroni-correction.*

| Participant | Comparison | Statistic (U) | P-value (raw) | P-value (adjusted) | Significance |
|---|---|---|---|---|---|
| P2 | Game vs Scroll | 4155 | 2.38E-02 | 7.15E-02 | ns |
| P2 | Reaction vs Game | 2776 | 4.28E-26 | 1.28E-25 | **** |
| P2 | Reaction vs Scroll | 191 | 9.29E-14 | 2.79E-13 | **** |
| P3 | Reaction vs Game | 2358 | 1.85E-06 | - | **** |
| C1 | Game vs Scroll | 4306 | 6.71E-17 | 2.01E-16 | **** |
| C1 | Reaction vs Game | 6911 | 1.09E-01 | 3.27E-01 | ns |
| C1 | Reaction vs Scroll | 1317 | 1.37E-15 | 4.10E-15 | **** |

***Table S16.*** *Welch's one-way ANOVA comparing mean z-scored force during the 200 ms following action onset across the cue-based (reaction time), competitive (Flip-that-Bucket), and unrestricted (scroll) tasks. Significance codes: ns, p ≥ .05; *p < .05; **p < .01; ***p < .001; ****p < .0001.*

| Comparison | Statistic (F) | DF1 | DF2 | P-value (raw) | Significance |
|---|---|---|---|---|---|
| Reaction time vs game vs scroll | 11.93 | 2 | 296.62 | 1.04E-05 | **** |

***Table S17.*** *Pairwise Welch's t-test results comparing mean z-scored force during the 200 ms following action onset across the cue-based (reaction time), competitive (Flip-that-Bucket), and unrestricted (scroll) tasks. P-values were Bonferroni-corrected across the three pairwise comparisons. Significance codes: ns, p ≥ .05; *p < .05; **p < .01; ***p < .001; ****p < .0001.*

| Task 1 | Task 2 | Task 1 mean | Task 2 mean | Diff. | Statistic (t) | DF | P-value (raw) | P-value (adjusted) | Sign. |
|---|---|---|---|---|---|---|---|---|---|
| Reaction time | Game | 1.76 | 1.77 | -0.01 | -0.13 | 285.45 | 8.99E-01 | 1.00E+00 | ns |
| Reaction time | Scroll | 1.76 | 1.39 | 0.38 | 3.65 | 329.55 | 3.08E-04 | 9.25E-04 | *** |
| Game | Scroll | 1.77 | 1.39 | 0.39 | 4.84 | 202.46 | 2.59E-06 | 7.77E-06 | **** |
